# Efficient production of saffron crocins and picrocrocin in *Nicotiana benthamiana* using a virus-driven system

**DOI:** 10.1101/2019.12.18.880765

**Authors:** Maricarmen Martí, Gianfranco Diretto, Verónica Aragonés, Sarah Frusciante, Oussama Ahrazem, Lourdes Gómez-Gómez, José-Antonio Daròs

## Abstract

Crocins and picrocrocin are glycosylated apocarotenoids responsible, respectively, for the color and the unique taste of the saffron spice, known as red gold due to its high price. Several studies have also shown the health-promoting properties of these compounds. However, their high costs hamper the wide use of these metabolites in the pharmaceutical sector. We have developed a virus-driven system to produce remarkable amounts of crocins and picrocrocin in adult *Nicotiana benthamiana* plants in only two weeks. The system consists of viral clones derived from tobacco etch potyvirus that express specific carotenoid cleavage dioxygenase (CCD) enzymes from *Crocus sativus* and *Buddleja davidii*. Metabolic analyses of infected tissues demonstrated that the sole virus-driven expression of *C. sativus* CsCCD2L or *B. davidii* BdCCD4.1 resulted in the production of crocins, picrocrocin and safranal. Using the recombinant virus that expressed CsCCD2L, accumulations of 0.2% of crocins and 0.8% of picrocrocin in leaf dry weight were reached in only two weeks. In an attempt to improve apocarotenoid content in *N. benthamiana*, co-expression of CsCCD2L with other carotenogenic enzymes, such as *Pantoea ananatis* phytoene synthase (PaCrtB) and saffron β-carotene hydroxylase 2 (BCH2), was performed using the same viral system. This combinatorial approach led to an additional crocin increase up to 0.35% in leaves in which CsCCD2L and PaCrtB were co-expressed. Considering that saffron apocarotenoids are costly harvested from flower stigma once a year, and that *Buddleja* spp. flowers accumulate lower amounts, this system may be an attractive alternative for the sustainable production of these appreciated metabolites.

## 1. Introduction

Plants possess an extremely rich secondary metabolism and, in addition to food, feed, fibers and fuel, also provide highly valuable metabolites for food, pharma and chemical industries. However, some of these metabolites are sometime produced in very limiting amounts. Plant metabolic engineering can address some of these natural limitations for nutritional improvement of foods or to create green factories that produce valuable compounds (Martin and Li, 2017; Yuan and Grotewold, 2015). Yet, the complex regulation of plant metabolic pathways and the particularly time-consuming approaches for stable genetic transformation of plant tissues highly limit quick progress in plant metabolic engineering. In this context, virus-derived vectors that fast and efficiently deliver biosynthetic enzymes and regulatory factors into adult plants may significantly contribute to solve some challenges (Sainsbury et al., 2012).

Carotenoids constitute an important group of natural products that humans cannot biosynthesize and must uptake from the diet. These compounds are synthesized by higher plants, algae, fungi and bacteria (Rodriguez-Concepcion et al., 2018). Yet, they play multiple roles in human physiology (Fiedor and Burda, 2014). Carotenoids are widely used as food colorants, nutraceuticals, animal feed, cosmetic additives and health supplements (Fraser and Bramley, 2004). In all living organisms, carotenoids act as substrates for the production of apocarotenoids (Ahrazem et al., 2016a). These are not only the simple breakdown products of carotenoids, as they act as hormones and are involved in signaling (Eroglu and Harrison, 2013; Jia et al., 2018; Walter et al., 2010). Apocarotenoids are present as volatiles, water soluble and insoluble compounds, and among those soluble, crocins are the most valuable pigments used in the food, and in less extent, in the pharmaceutical industries (Ahrazem et al., 2015). Crocins are glycosylated derivatives of the apocarotenoid crocetin. They are highly soluble in water and exhibit a strong coloring capacity. In addition, crocins are powerful free radical quenchers, a property associated to the broad range of health benefits they exhibit (Bukhari et al., 2018; Christodoulou et al., 2015; Georgiadou et al., 2012; Nam et al., 2010). The interest in the therapeutic properties of these compounds is increasing due to their analgesic and sedative properties (Amin and Hosseinzadeh, 2012), and neurological protection and anticancer activities (Finley and Gao, 2017; Skladnev and Johnstone, 2017). Further, clinical trials indicate that crocins have a positive effect in the treatment of depression and dementia (Lopresti and Drummond, 2014; Mazidi et al., 2016). Other apocarotenoids, such as safranal (2,6,6-trimethyl-1,3-cyclohexadiene-1-carboxaldehyde) and its precursor picrocrocin (β-D-glucopyranoside of hydroxyl-β-cyclocitral), have also been shown to reduce the proliferation of different human carcinoma cells (Cheriyamundath et al., 2018; Jabini et al., 2017; Kyriakoudi et al., 2015) and to exert anti-inflammatory effects (Zhang et al., 2015).

*Crocus sativus* L. is the main natural source of crocins and picrocrocin, respectively responsible for the color and flavor of the highly appreciated spice, saffron (Tarantilis et al., 1995). Crocin pigments accumulate at huge levels in the flower stigma of these plants conferring this organ with a distinctive dark red coloration (Moraga et al., 2009). Yet, these metabolites reach huge prices in the market, due to the labor intensive activities associated to harvesting and processing the stigma collected from *C. sativus* flowers (Ahrazem et al., 2015). Besides *C. sativus*, gardenia (*Gardenia* spp.) fruits are also a commercial source of crocins, but at much lower scale. In addition, gardenia fruits do not accumulate picrocrocin (Moras et al., 2018; Pfister et al., 1996). Some other plants, such as *Buddleja* spp., also produce crocins, although they are not commercially exploited due to low accumulation (Liao et al., 1999). All these plants share the common feature of expressing carotenoid cleavage dioxygenase (CDD) activities, like the saffron CCD2L or the CCD4 subfamily recently identified in *Buddleja* spp. (Ahrazem et al., 2017).

In *C. sativus* and *Buddleja* spp., zeaxanthin is the precursor of crocetin (Fig. 1). Cleavage of the zeaxanthin molecule at the 7,8 and 7′,8′ double bonds render one molecule of crocetin dialdehyde and two molecules of 4-hydroxy-2,6,6-trimethyl-1-cyclohexene-1-carboxaldehyde (HTCC) (Ahrazem et al., 2017, 2016c; Frusciante et al., 2014). Crocetin dialdehyde is further transformed to crocetin by the action of aldehyde dehydrogenase (ALDH) enzymes (Demurtas et al., 2018; Gómez-Gómez et al., 2018, 2017). Next, crocetin serves as substrate of glucosyltransferase (UGT) activities that catalyze the production of crocins through transfer of glucose to both ends of the molecule (Côté et al., 2001; Demurtas et al., 2018; Moraga et al., 2004; Nagatoshi et al., 2012). The HTCC molecule is also recognized by UGTs, resulting in production of picrocrocin (Fig. 1) (Diretto et al., 2019a). In *C. sativus*, the plastidic CsCCD2L enzyme catalyzes the cleavage of zeaxanthin at 7,8;7′,8′ double bonds (Ahrazem et al., 2016c; Frusciante et al., 2014), which is closely related to the CCD1 subfamily (Ahrazem et al., 2016b, 2016a). In *Buddleja davidii*, the CCD enzymes that catalyze the same reaction belong to a novel group within the CCD4 subfamily of CCDs (Ahrazem et al., 2016a). These *B. davidii* enzymes, BdCCD4.1 and BdCCD4.3, also localize in plastids and are expressed in the flower tissue (Ahrazem et al., 2017).

**Fig. 1.**
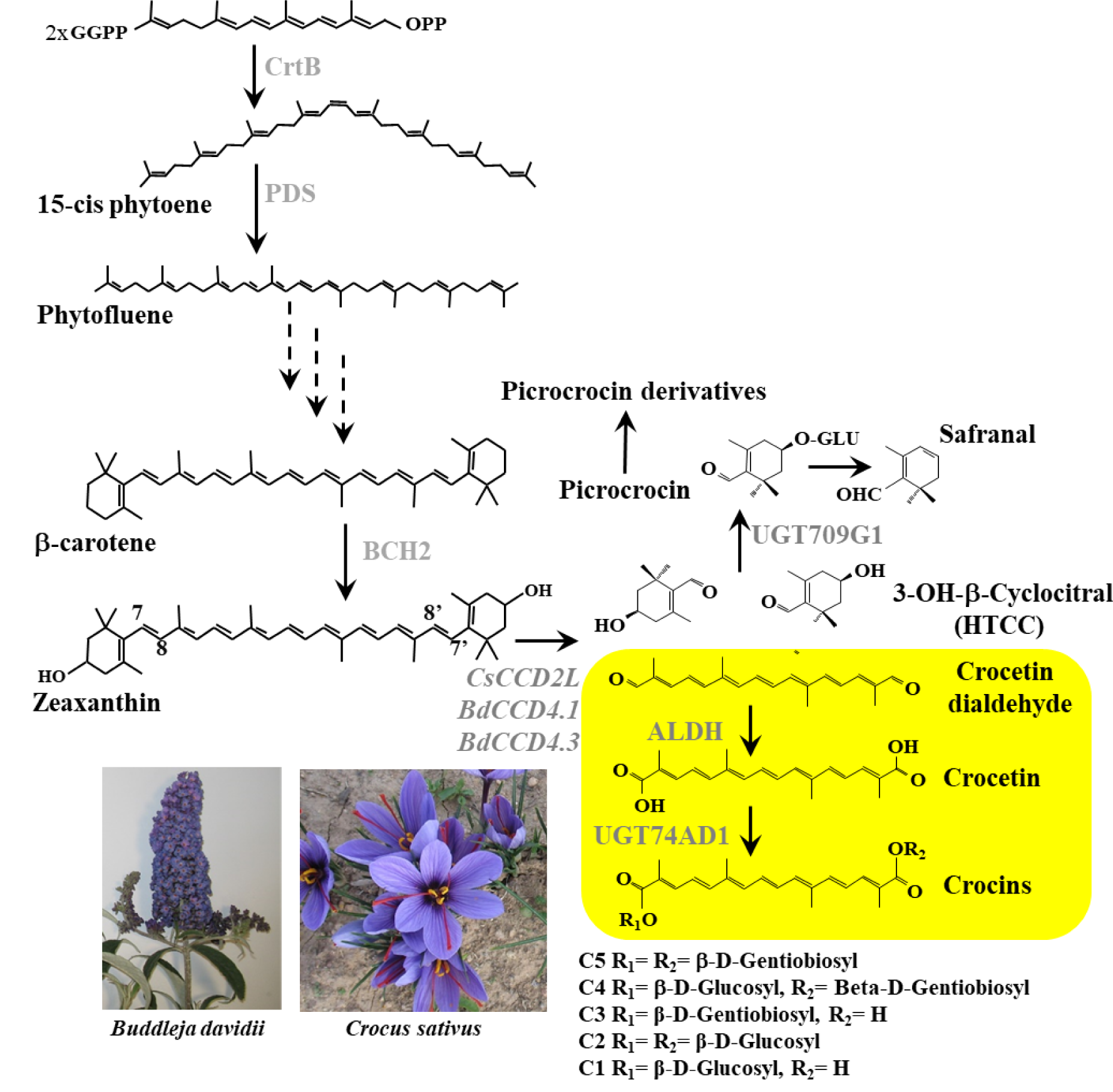
Schematic overview of the crocins biosynthesis pathway in *C. sativus* and *B. davidii*. CrtB, phytoene synthase; PDS, phytoene desaturase; BCH2, carotene hydroxylase; CsCCD2L, *C. sativus* carotenoid cleavage dioxygenase 2L; BdCCD4.1 and 4.3, *B. davidii* carotenoid cleavage dioxygenase 4.1 and 4.3; ALDH, aldehyde dehydrogenase; UGT74AD1, UDP-glucosyltransferase 74AD1.

Due to its high scientific and commercial interest, crocetin and crocins biosynthesis has arisen great attention, and attempts to metabolically engineer their pathway in microbial systems have been reported, although with limited success (Chai et al., 2017; Diretto et al., 2019a; Tan et al., 2019; Wang et al., 2019). More in detail, the use of the saffron CCD2L enzyme resulted in a maximum accumulation of 1.22 mg/l, 15.70 mg/l, and 4.42 mg/l of crocetin in, respectively, *Saccharomyces cerevisiae* (Chai et al., 2017; Tan et al., 2019) and *Escherichia coli* (Wang et al., 2019). In a subsequent study aimed to confirm, *in planta*, the role of a novel UGT in picrocrocin biosynthesis (Diretto et al., 2019a), *Nicotiana benthamiana* leaves were transiently transformed, *via Agrobacterium tumefaciens*, with CCD2L, alone or in combination with UGT709G1, which led to the production of 30.5 μg/g DW of crocins, with a glycosylation degree ranging from 1 to 4 (crocin 1-4) (unpublished data). Although promising, these results cannot be considered efficient and sustainable in the context of the development of an industrial system for the production of the saffron high-value apocarotenoids.

Here we used a viral vector derived from *Tobacco etch virus* (TEV, genus *Potyvirus*; family *Potyviridae*) (Bedoya et al., 2010) to transiently express a series of CCD enzymes, alone or in combination with other carotenoid and non-carotenoid biosynthetic enzymes, in *N. benthamiana* plants. Interestingly, tissues infected with viruses that expressed CsCCD2L or BdCCD4.1 acquired yellow pigmentation visible to the naked eye. Metabolite analyses of these tissues demonstrated the accumulation of crocins and picrocrocin, reaching levels attractive enough for their commercial production, up to 0.35% and 0.8% of DW, respectively, in leaves in which CsCCD2L and *Pantoea ananatis* phytoene synthase (PaCrtB) were simultaneously expressed.

## 2. Materials and Methods

### 2.1. Plasmid construction

Plasmids were constructed by standard molecular biology techniques, including PCR amplifications of cDNAs with the high-fidelity Phusion DNA polymerase (Thermo Scientific) and Gibson DNA assembly (Gibson et al., 2009), using the NEBuilder HiFi DNA Assembly Master Mix (New England Biolabs). To generate a transformed *N. benthamiana* line that stably expresses TEV NIb, we built the plasmid p235KNIbd. This plasmid is a derivative of pCLEAN-G181 (Thole et al., 2007) and between the left and right borders of *A. tumefaciens* transfer DNA (T-DNA) contains two in tandem cassettes in which the neomycin phosphotransferase (NPTII; kanamycin resistance) and the TEV NIb are expressed under the control of CaMV 35S promoter and terminator. The exact nucleotide sequence of the p235KNIbd T-DNA is in Supplementary Fig. S1. Plasmids pGTEVΔNIb-CsCCD2L, -BdCCD4.1, -BdCCD4.3, -CsCCD2L/PaCrtB, - CsCCD2L/CsBCH2 and -CsCCD2L/CsLPT1 were built on the basis of pGTEVa (Bedoya et al., 2012) that contains a wild-type TEV cDNA (GenBank accession number DQ986288 with the two silent and neutral mutations G273A and A1119G) flanked by the CaMV 35S promoter and terminator in a binary vector that also derives from pCLEAN-G181. These plasmids were built, as explained above, by PCR amplification of CsCCD2L (Ahrazem et al., 2016c), BdCCD4.1 and BdCCD4.3 (Ahrazem et al., 2017), PaCrtB (Majer et al., 2017), CsBCH2 (Castillo et al., 2005) and CsLPT1 (Gómez-Gómez et al., 2010) cDNAs and assembly into a pGTEVa version in which the NIb cistron was deleted (pGTEVaΔNIb) (Majer et al., 2015). The exact sequences of the TEV recombinant clones (TEVΔNIb-CsCCD2L, -BdCCD4.1, -BdCCD4.3, -CsCCD2L/PaCrtB, - CsCCD2L/CsBCH2 and -CsCCD2L/CsLPT1) contained in the resulting plasmids are in Supplementary Fig. S2. The plasmid expressing the recombinant TEVΔNIb-aGFP clone (Supplementary Fig. S2) was previously described (Majer et al., 2015).

### 2.2. Plant transformation

The LBA4404 strain of *A. tumefaciens* was sequentially transformed with the helper plasmid pSoup (Hellens et al., 2000) and p235KNIbd. Clones containing both plasmids were selected in plates containing 50 µg/ml rifampicin, 50 µg/ml kanamycin and 7.5 µg/ml tetracycline. A liquid culture grown from a selected colony was used to transform *N. benthamiana* (Clemente, 2006). Briefly, *N. benthamiana* leaf explants were incubated with the *A. tumefaciens* culture at an optical density (600 nm) of 0.5 for 30 min and then incubated on solid media. Four days later, explants were transferred to plates with organogenesis media containing 100 µg/ml of kanamycin. Explants were repeatedly transferred to fresh organogenesis plates every three weeks until shoots emerged. These shoots were first transferred to rooting media and, when roots appeared, cultivated in a greenhouse with regular soil until seeds were harvested. Transformation with the *NIb* cDNA was confirmed by PCR. Batches of seedlings from four independent transformed lines were screened for efficient complementation of TEV clones lacking NIb by mechanical inoculation of TEVΔNIb-Ros1 (Bedoya et al., 2010).

### 2.3. Plant inoculation

*A. tumefaciens* C58C1 competent cells, previously transformed with the helper plasmid pCLEAN-S48 (Thole et al., 2007), were electroporated with different plasmids that contained the TEV recombinant clones and selected in plates with 50 µg/ml rifampicin, 50 µg/ml kanamycin and 7.5 µg/ml tetracycline. Liquid cultures of selected colonies were brought to an optical density of 0.5 (600 nm) in agroinoculation solution (10 mM MES-NaOH, pH 5.6, 10 mM MgCl_2_ and 150 μM acetosyringone) and incubated for 2 h at 28°C. Using a needleless syringe, these cultures were used to infiltrate one leaf of five-week-old plants of the selected *N. benthamiana* transformed line that stably expresses TEV NIb. After inoculation, plants were kept in a growth chamber at 25°C under a 12 h day-night photoperiod. Leaves were collected at different dpi as indicated; in the case of the time-course experiments, leaf samples were collected at 8 time-points (0, 4, 8, 11, 13, 15, 18 and 20 dpi).

### 2.4. Analysis of virus progeny

Viral progeny was analyzed by reverse transcription (RT)-PCR using RevertAid reverse transcriptase (Thermo Scientific) and Phusion DNA polymerase followed by electrophoretic separation of the amplification products in 1% agarose gels that were stained with ethidium bromide. RNA from *N. benthamiana* plants was purified from upper non-inoculated leaves at 15 dpi using silica-gel columns (Zymo Research). For virus infection diagnosis, a cDNA corresponding to the TEV CP cistron (804 bp) was amplified. Aliquots of the RNA preparations were subjected to RT using primer PI (5’-CTCGCACTACATAGGAGAATTAGAC-3’), followed by PCR amplification with primers PII (5’-AGTGGCACTGTGGGTGCTGGTGTTG-3’) and PIII (5’-CTGGCGGACCCCTAATAG-3’). To analyze the potential deletion of the inserted CDD cDNAs, RNA aliquots were reverse transcribed using primer PIV (5’-GCTGTTTGTCACTCAATGACACATTAT-3’) and amplified by PCR with primers PV (5’-AAAATAACAAATCTCAACACAACATATAC-3’) and PVI (5’-CCGCGGTCTCCCCATTATGCACAAGTTGAGTGGTAGC-3’).

### 2.5. Metabolite extraction and analysis

Polar and apolar metabolites were extracted from 50 mg of lyophilized leaf tissue. For polar metabolites analyses (crocins and picrocrocin), the tissues were extracted in cold 50% methanol. The soluble fraction was analyzed by HPLC-DAD-HRMS and HPLC-DAD (Ahrazem et al., 2018; Moraga et al., 2009). The liposoluble fractions (crocetin, HTCC, -cyclocitral, carotenoids and chlorophylls) were extracted with 0.5:1 ml cold extraction solvents (50:50 methanol and CHCl_3_), and analyzed by HPLC-DAD-HRMS and HPLC-DAD as previously described (Ahrazem et al., 2018; Castillo et al., 2005; D’Esposito et al., 2017; Fasano et al., 2016). Metabolites were identified on the basis of absorption spectra and retention times relative to standard compounds. Pigments were quantified by integrating peak areas that were converted to concentrations in comparison with authentic standards. Chlorophyll a and b were determined by measuring absorbance at 645 nm and 663 nm as previously described (Richardson et al., 2002). Mass spectrometry carotenoid, chlorophyll derivative and apocarotenoid identification and quantification was carried out as previously described (Ahrazem et al., 2018; Diretto et al., 2019a; Rambla et al., 2016). Mass spectrometry analysis of other isoprenoids (tocochromanols and quinones) were performed as reported before (Sulli et al., 2017). Picrocrocin derivatives, as reported by (Moras et al., 2018) and (D’Archivio et al., 2016), were tentatively identified according to their m/z accurate masses as included in the PubChem database for monoisotopic masses or by using the Mass Spectrometry Adduct Calculator from the Metabolomics Fiehn Lab for adduct ions; and were further validated by isotopic pattern ratio and the comparison between theoretical and experimental m/z fragmentation, by using the MassFrontier 7.0 software (Thermo Fisher Scientific).

### 2.6. Subcellular localization of crocins

To determine the subcellular localization of crocins in *N. benthamiana* tissues, LSCM was performed with a Zeiss 7080 Axio Observer equipped with a water immersion objective lens (C-Apochromat 40X/1.20 W). An argon ion laser at 458 nm was used for excitation. Crocins and chloroplast autofluorescence were detected with 520 to 540 nm and 660 to 750 nm band-pass emission filters, respectively. Images were processed using the FIJI software (http://fiji.sc/Fiji).

### 2.7. Statistics and bioinformatics

Statistical validation of the data (ANOVA+ Tukey’s t-test) and heatmap visualization were performed as previously described (Cappelli et al., 2018; Grosso et al., 2018).

## 3. Results

### 3.1. TEV recombinant clones that express CCD enzymes in N. benthamiana

The goal of this work was to develop a heterologous system to efficiently produce highly appreciated and scarce apocarotenoids in plant tissues. For this purpose, we planned to use an expression vector derived from TEV, more specifically TEVΔNIb (Bedoya *et al*., 2010), recently employed to rewire the lycopene biosynthetic pathway from the plastid to the cytosol of plant cells (Majer et al., 2017). Since CCD is the first enzyme in the apocarotenoid biosynthetic pathway (Fig. 1), we started exploring the virus-based expression of a series of CCD enzymes from well-known crocins producing plants: namely *C. sativus* CsCCD2L (Ahrazem et al., 2016c), and *B. davidii* BdCCD4.1 and BdCCD4.3 (Ahrazem et al., 2017).

In TEVΔNIb, the approximately 1.6 kb corresponding to nuclear inclusion *b* (NIb) cistron, coding for the viral RNA-dependent RNA polymerase, is deleted to increase the space to insert foreign genetic information. Then, the viral recombinant clones only infect plants in which NIb is supplied in *trans* (Bedoya et al., 2010). We chose *N. benthamiana* as the biofactory plant for apocarotenoid production because of the amenable genetic transformation, the high amounts of biomass achievable and the fast growth. For this aim, as the first step in our work, we transformed *N. benthamiana* leaf tissue using *Agrobacterium tumefaciens* and regenerated adult plants that constitutively expressed TEV NIb under the control of *Cauliflower mosaic virus* (CaMV) 35S promoter and terminator. To screen for transformed lines that efficiently complemented TEVΔNIb deletion mutants, we inoculated the plants with TEVΔNIb-Ros1, a viral clone in which the NIb cistron was replaced by a cDNA encoding for the MYB-type transcription factor Rosea1 from *Antirrhinum majus* L., whose expression induces accumulation of colored anthocyanins in infected tissues and facilitates visual tracking of infection (Bedoya et al., 2010, 2012). On the basis of anthocyanin accumulation, we selected a *N. benthamiana* transformed line that efficiently complemented the replication and systemic movement of TEVΔNIb-Ros1 (Supplementary Fig. S3). Then, by self-pollination and TEVΔNIb-Ros1 inoculation of the progeny, we selected a transformed homozygous line that was used in all subsequent experiments.

Next, on the context of a TEV lacking NIb, we constructed three recombinant clones (Fig. 2A) to express *C. sativus CsCCD2L* (TEVΔNIb-CsCCD2L), *B. davidii BdCCD4.1* and *BdCCD4.3* (TEVΔNIb-BdCCD4.1 and TEVΔNIb-BdCCD4.3). These CCDs are plastidic enzymes that are encoded in the nucleus and contain native amino-terminal transit peptides. To target these enzymes to plastids, we inserted their cDNAs in a position corresponding to the amino terminus of the viral polyprotein in the virus genome (Majer et al., 2015). CCD cDNAs also contained a sequence corresponding to an artificial NIaPro cleavage site at the 3’ end to mediate the release of the heterologous protein from the viral polyprotein. Based on previous observations, we used the −8/+3 site that splits NIb and CP in TEV. The exact sequences of these clones are in Supplementary Fig. S2.

**Fig. 2.**
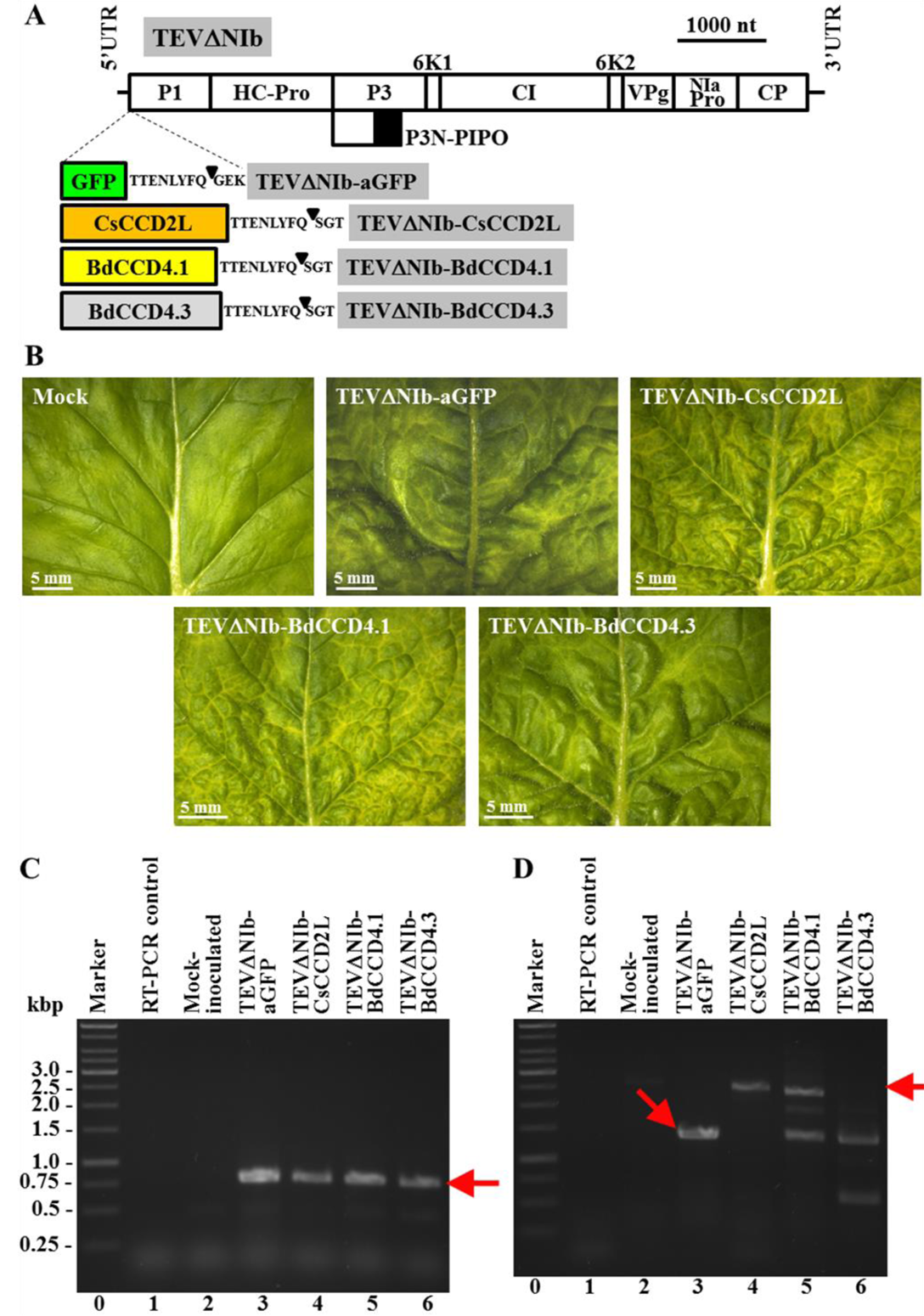
Inoculation of *N. benthamiana* plants that stably express NIb with TEVΔNIb recombinant clones that express different CCDs. (A) Schematic representation of TEVΔNIb genome indicating the position where the GFP (green box) and the different CCDs (CsCCD2L, orange box; BdCCD4.1, yellow box; and BdCCD4.3, gray box) were inserted. The sequence of the artificial NIaPro cleavage site to mediate the release of the recombinant proteins from the viral polyprotein is also indicated. The black triangle indicates the exact cleavage site. Lines represent TEV 5’ and 3’ UTR and boxes represent P1, HC-Pro, P3, P3N-PIPO, 6K1, CI, 6K2, VPg, NIaPro and CP cistrons, as indicated. Scale bar corresponds to 1000 nt. (B) Pictures of representative leaves from plants mock-inoculated and agroinoculated with TEVΔNIb-aGFP, TEVΔNIb-CsCCD2L, TEVΔNIb-BdCCD4.1 and TEVΔNIb-BdCCD4.3, as indicated, taken at 13 dpi. Scale bars correspond to 5 mm. (C) Virus diagnosis and (D) analysis of the progeny of recombinant TEV in *N. benthamiana* plants mock-inoculated and agroinoculated with TEVΔNIb-aGFP, TEVΔNIb-CsCCD2L, -BdCCD4.1 and -BdCCD4.3. RNA was extracted at 15 dpi and subjected to RT-PCR amplification. PCR products were separated in a 1% agarose gel that was stained with ethidium bromide. Representative samples of triplicate analyses are shown. (C and D) Lanes 0, DNA marker ladder with sizes (in bp) on the left; lanes 1, RT-PCR controls with no RNA added; lanes 2, mock-inoculated plants; lanes 3 to 6, plants inoculated with TEVΔNIb-aGFP (lanes 3), TEVΔNIb-CsCCD2L (lanes 4), TEVΔNIb-BdCCD4.1 (lanes 4) and TEVΔNIb-BdCCD4.3 (lanes 5). The arrows point to the bands corresponding to (C) the TEV CP, and (D) the full-length viral progenies cDNAs.

Finally, transformed *N. benthamiana* plants that express NIb were agroinoculated with the three TEV recombinant clones. As a control in this experiment, some plants were agroinoculated with TEVΔNIb-aGFP (Fig. 2A and Supplementary Fig. S2), which is a recombinant clone that expresses GFP from the amino-terminal position in the viral polyprotein. Interestingly, 7 days post-inoculation (dpi), even before symptoms were observed, we noticed a distinctive yellow pigmentation in systemic tissues of the plants agroinoculated with TEVΔNIb-CsCCD2L. Symptoms of infection were soon observed (approximately 8 dpi) in plants agroinoculated with TEVΔNIb-aGFP,-CsCCD2L and -BdCCD4.1. Distinctive yellow pigmentation was also observed in symptomatic tissues of plants agroinoculated with TEVΔNIb-BdCCD4.1, although with some delay comparing to TEVΔNIb-CsCCD2L. Yellow pigmentation of symptomatic tissues in plants inoculated with TEVΔNIb-CsCCD2L was more intense than that in tissues of plants inoculated with TEVΔNIb-BdCCD4.1. Plants inoculated with TEVΔNIb-BdCCD4.3 showed mild symptoms of infection at approximately 13 dpi and yellow pigmentation was never observed. Fig. 2B shows pictures of representative leaves of all these plants at 13 dpi. Reverse transcription (RT)-PCR analysis confirmed TEV infection in all inoculated plants (Fig. 2C). Observation of symptomatic tissues with a fluorescence stereomicroscope confirmed expression of GFP only in plants inoculated with the TEVΔNIb-aGFP control (Supplementary Fig. S4). Analysis of viral progeny at 15 dpi by RT-PCR indicated that, while the inserts of TEVΔNIb-aGFP, -CsCCD2L are perfectly stable in the recombinant virus, those of TEVΔNIb-BdCCD4.1 and TEVΔNIb-BdCCD4.3 are partially and completely lost, respectively, at this time post-inoculation (Fig. 2D). Based on distinctive yellow color, these results suggested that the virus-driven expression of CsCCD2L or BdCCD4.1 in *N. benthamiana* tissues may induce apocarotenoid accumulation.

### 3.2. Analysis of apocarotenoids in infected tissues

Symptomatic leaf tissues from plants infected with TEVΔNIb-aGFP, TEVΔNIb-CsCCD2L and TEVΔNIb-BdCCD4.1 were subjected to extraction and analysis by high performance liquid chromatography-diode array detector-high resolution mass spectrometry (HPLC-DAD-HRMS) to determine their apocarotenoid and carotenoid profiles. Tissues from mock-inoculated controls were also included in the analysis. The tissues from the plants infected with TEVΔNIb-BdCCD4.3 were discarded due to the absence of pigmentation and rapid deletion of BdCCD4.3 cDNA in the viral progeny. Analysis of the polar fraction of tissues infected with TEVΔNIb-CsCCD2L and TEVΔNIb-BdCCD4.1 showed a series of peaks with maximum absorbance from 433 to 439 nm. These peaks were not observed in extracts from tissues from mock-inoculated plants or plants infected with the TEVΔNIb-aGFP control (Fig. 3). Mass spectrometry chromatograms evidenced the presence, for all the fore mentioned peaks, of the crocetin aglycon (m/z 329.1747 (M+H)), thus supporting the presence of crocins (Supplementary Fig. S5). Further analyses of the sugar moieties conjugated with crocetin lead to the identification of crocins with different degrees of glycosylation from one glucose molecule to five in the infected tissues in which CCDs were expressed (Fig. 4A and Supplementary Table S1). However, predominant crocins were those conjugated with three and four glucose molecules. Interestingly, CCD-infected *N. benthamiana* leaves, displayed a different pattern of crocin accumulation compared to that typical from saffron stigma (Fig. 3A), with trans-crocin 4, followed by trans-crocin 3 and 2, being the most abundant crocin species. Minor changes were also observed between TEVΔNIb-CsCCD2L and TEVΔNIb-BdCCD4.1 samples (Supplementary Table S2): in the former, trans-crocin 3 was the most abundant (30.84%), followed by cis-crocin 4 (17.04%) and trans-crocin 2 (16.84%); whereas similar amounts in cis-crocin 5, cis-crocin 4 and trans-crocin 3 were found in the latter (20.92, 20.62 and 18.31%, respectively). Picrocrocin, safranal and several derivatives were also identified in the polar fraction of tissues infected with TEVΔNIb-CsCCD2L and TEVΔNIb-BdCCD4.1 (Fig. 4B and Supplementary Table S1). All these polar apocarotenoids were absent in the control tissues non-infected and infected with TEVΔNIb-aGFP (Fig. 4B). Overall, and in agreement with the most intense visual phenotype, TEVΔNIb-CsCCD2L-infected tissues displayed a higher accumulation in all the linear and cyclic apocarotenoids than TEVΔNIb-BdCCD4.1-infected tissues (Fig. 4, Table 1 and Supplementary Table S1).

**Fig. 3.**
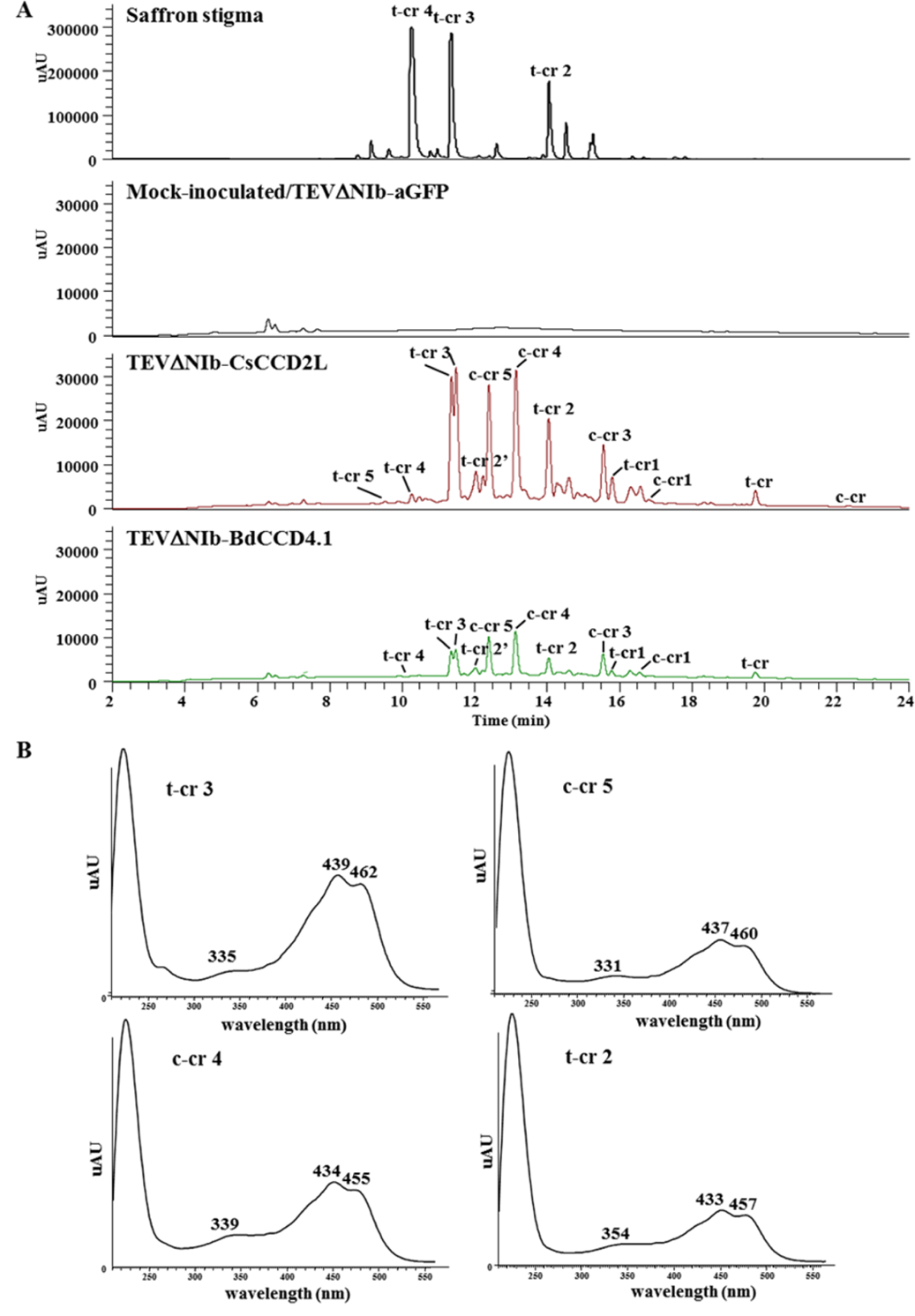
Heterologous production of crocins in *N. benthamiana* tissues infected with recombinant viruses expressing CCD enzymes. (A) Chromatographic profile run on an HPLC-PDA-HRMS and detected at 440 nm of the polar fraction of tissues infected with TEVΔNIb-aGFP, TEVΔNIb-CsCCD2L and TEVΔNIb-BdCCD4.1. The profile of saffron stigma is also included as a control. Peaks abbreviations correspond to: c-cr, *cis*-crocetin; t-cr, *trans*-crocetin; c-cr1, *cis*-crocin 1; t-cr1, *trans*-crocin 1; t-cr2, *trans*-crocin 2; t-cr2ʹ, *trans*-crocin 2ʹ; c-cr3, *cis*-crocin 3, t-cr3, *trans*-crocin 3; c-cr4, *cis*-crocin 4; t-cr4, *trans*-crocin 4; c-cr5, *cis*-crocin 5; t-cr5, *trans*-crocetin 5. (B) Absorbance spectra of the major crocins detected in the polar extracts of *N. benthamiana* tissues infected with TEVΔNIb-CsCCD2L and TEVΔNIb-BdCCD4.1. Analyses were performed at 13 dpi.

**Fig. 4.**
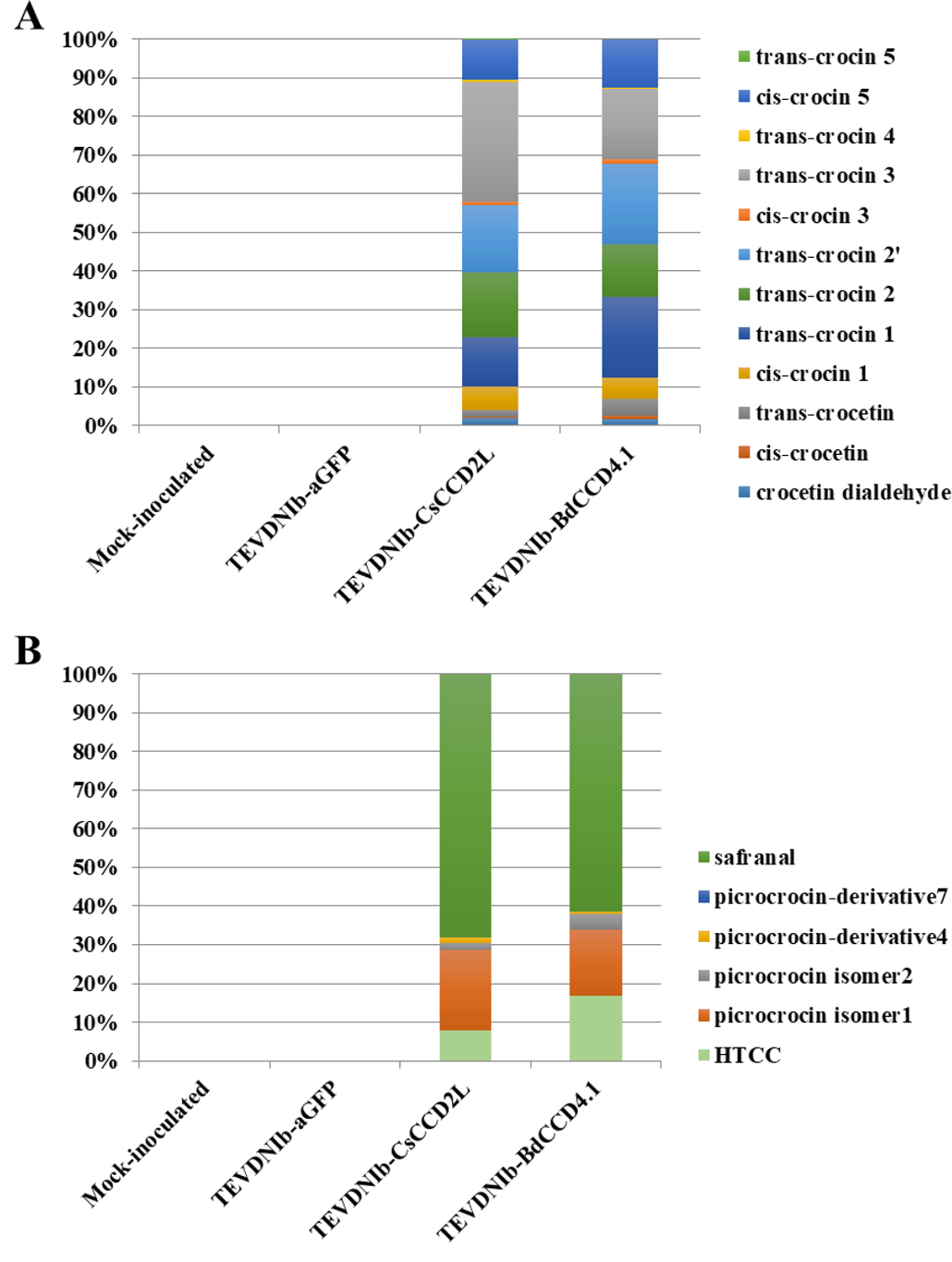
Apocarotenoid content and composition of *N. benthamiana* tissues from mock-inoculated plants and plants infected with TEVΔNIb-aGFP, TEVΔNIb-CsCCD2L and TEVΔNIb-BdCCD4.1. (A) Crocetin dialdehyde, crocetin and crocins accumulation. (B) Levels of safranal, picrocrocin and its derivatives, and HTCC. Data are averages ± sd of three biological replicates and expressed as % of fold internal standard (IS) levels. Analyses were performed at 13 dpi.

**Table 1.** Ratios of (A) crocins- and (B) picrocrocin-related apocarotenoid levels in tissues infected with TEVΔNIb-CsCCD2L and TEVΔNIb-BdCCD4.1.

| **A** | **CsCCD2L/BdCCD4.1** |
| --- | --- |
| crocetin dialdehyde | 3.206 |
| cis-crocetin | 1.139 |
| trans-crocetin | 1.113 |
| cis-crocin 1 | 3.318 |
| trans-crocin 1 | 1.755 |
| trans-crocin 2 | 3.605 |
| trans-crocin 2' | 2.420 |
| cis-crocin 3 | 2.137 |
| trans-crocin 3 | 4.915 |
| trans-crocin 4 | 5.580 |
| cis-crocin 5 | 2.448 |
| trans-crocin 5 | - |
| **B** | **CsCCD2L/BdCCD4.1** |
| HTCC | 1.573 |
| picrocrocin isomer1 | 4.237 |
| picrocrocin isomer2 | 1.405 |
| picrocrocin-derivivative4 | 8.577 |
| picrocrocin-derivivative7 | 6.474 |
| safranal | 3.805 |

In the polar and apolar fractions of TEVΔNIb-CsCCD2L and TEVΔNIb-BdCCD4.1-infected leaves, we also detected the presence of crocetin dialdehyde and HTCC, which are the products of the enzymatic cleavage step, and crocetin and safranal, which are produced by the activity of ALDH enzymes and the spontaneous deglycosilation of picrocrocin, respectively (Fig. 4 and Supplementary Table S1). None of these metabolites were found in the TEVΔNIb-aGFP-infected and mock-inoculated control tissues. Subsequently, the levels of different carotenoids and chlorophylls were also investigated in the apolar fractions (Fig. 5 and Supplementary Table S3). As expected, virus infection negatively affected the isoprenoid pools, with most of carotenoids and chlorophylls reduced in all infected tissues compared to those from mock-inoculated plants. However, the comparison between tissues infected with TEVΔNIb-aGFP and both CCD viruses highlighted a series of alterations at metabolite level. The most striking differences were in the contents of phytoene and phytofluene, as well as zeaxanthin and lutein. While the formers increased in the TEVΔNIb-CsCCD2L and TEVΔNIb-BdCCD4.1-infected tissues, the levels of zeaxanthin and lutein were strongly reduced (Fig. 5A and Table S3A). On the contrary, no significant alterations in chlorophyll levels were observed in tissues infected with the two CCD viruses when compared to the TEVΔNIb-aGFP-infected control (Fig. 5B and Supplementary Table S3B). Other typical leaf isoprenoids, such as tocochromanols and quinones, displayed only few changes in CCD versus control tissues: for instance, β-/γ-tocopherol, α-tocopherol quinone and menaquinone-8 increased, whereas plastoquinone was reduced in CsCCD2L and BdCCD4.1-infected tissues compared to mock-inoculated and GFP-infected leaves (Fig. 5C and Supplementary Table S3C).

**Fig. 5.**
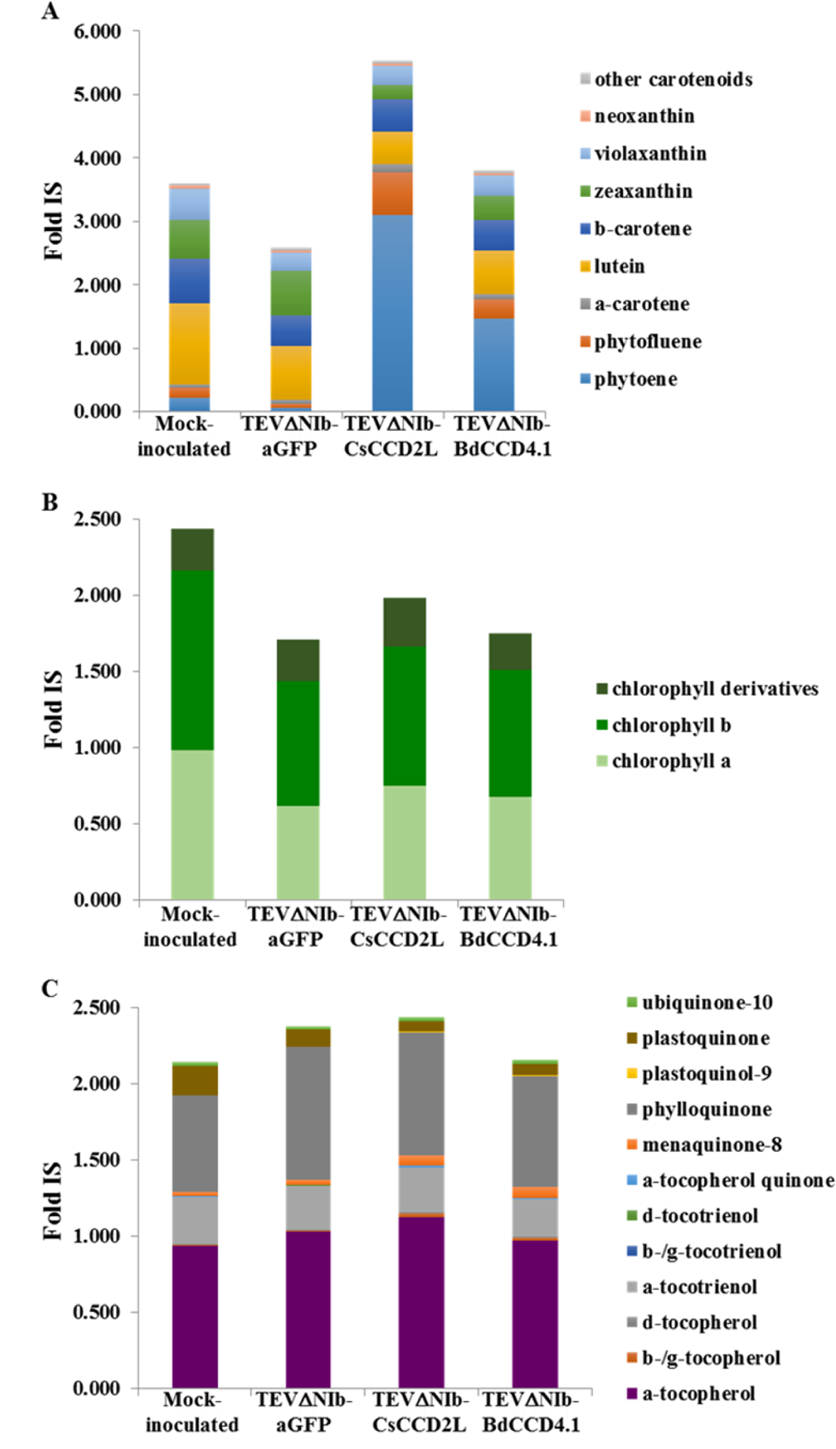
Relative quantities of (A) carotenoids (B) chlorophylls and (C) quinones and tocochromanols detected by HPLC-DAD-HRMS in tissues of mock-inoculated and infected (TEVΔNIb-aGFP, TEVΔNIb-CsCCD2L and TEVΔNIb-BdCCD4.1) *N. benthamiana* plants. Data are averages ± sd of three biological replicates and expressed as fold internal standard (IS). Analyses were performed at 13 dpi.

### 3.3. Subcellular accumulation of apocarotenoids in tissues infected with CCD-expressing viruses

On the basis of the differential autofluorescence of chlorophylls and carotenoids, we analyzed the subcellular localization of crocins that accumulate in symptomatic tissues of plants infected with TEVΔNIb-CsCCD2L and TEVΔNIb-BdCCD4.1. Laser scanning confocal microscopy (LSCM) images showed an intense fluorescence signal from crocins in the vacuoles, while tissues infected with TEVΔNIb only showed chlorophyll fluorescence (Fig. 6).

**Fig. 6.**
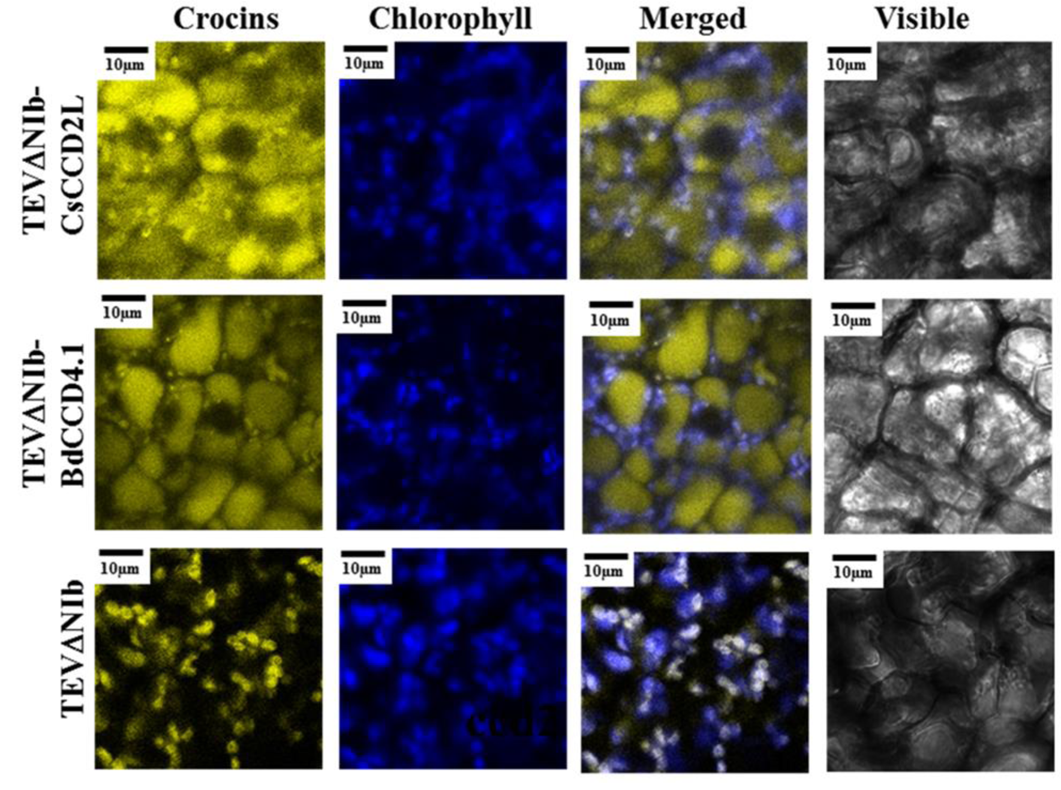
LSCM images of crocins and chlorophyll autofluorescence in symptomatic leaves of *N. benthamiana* plants infected with TEVΔNIb, TEVΔNIb-CsCCD2L and TEVΔNIb-BdCCD4.1, as indicated, at 13 dpi. Merged images of both fluorescent signals and images under white light are also shown. Scale bars indicate 10 μm.

### 3.4. Time course accumulation of crocins and picrocrocin in inoculated plants

Next, we researched the point with the maximum apocarotenoid accumulation in inoculated plants. For this aim, we carried out a time-course analysis focusing only on the virus recombinant clone TEVΔNIb-CsCCD2L that induces the highest apocarotenoid accumulation in *N. benthamiana* tissues. After plant agroinoculation, upper systemic tissues were collected at 0, 4, 8, 11, 13, 15, 18 and 20 dpi from three independent replicate plants per time point. Pigments were extracted and analyzed by HPLC-DAD. In these tissues, accumulation of total crocins increased from approximately 8 to 13 dpi, reaching a plateau up to the end of the analysis (20 dpi) (Fig. 7A). Picrocrocin levels showed identical behavior (Fig. 7B). Accumulation of lutein, β-carotene and chlorophylls was also evaluated in all these samples. In contrast to apocarotenoids, accumulation of lutein and β-carotene (Fig. 7C) as well as chlorophylls a and b (Fig. 7D) dropped as infection progressed. These results indicate that the virus-driven system to produce apocarotenoids in *N. benthamiana* developed here reaches the highest yield in only 13 days after inoculating the plants and this yield is maintained during at least one week with no apparent loss. This experiment also revealed the remarkable accumulation of 2.18±0.23 mg of crocins and 8.24±2.93 mg of picrocrocin per gram of *N. benthamiana* dry weight leaf tissue with the sole virus-driven expression of *C. sativus* CsCCD2L.

**Fig. 7.**
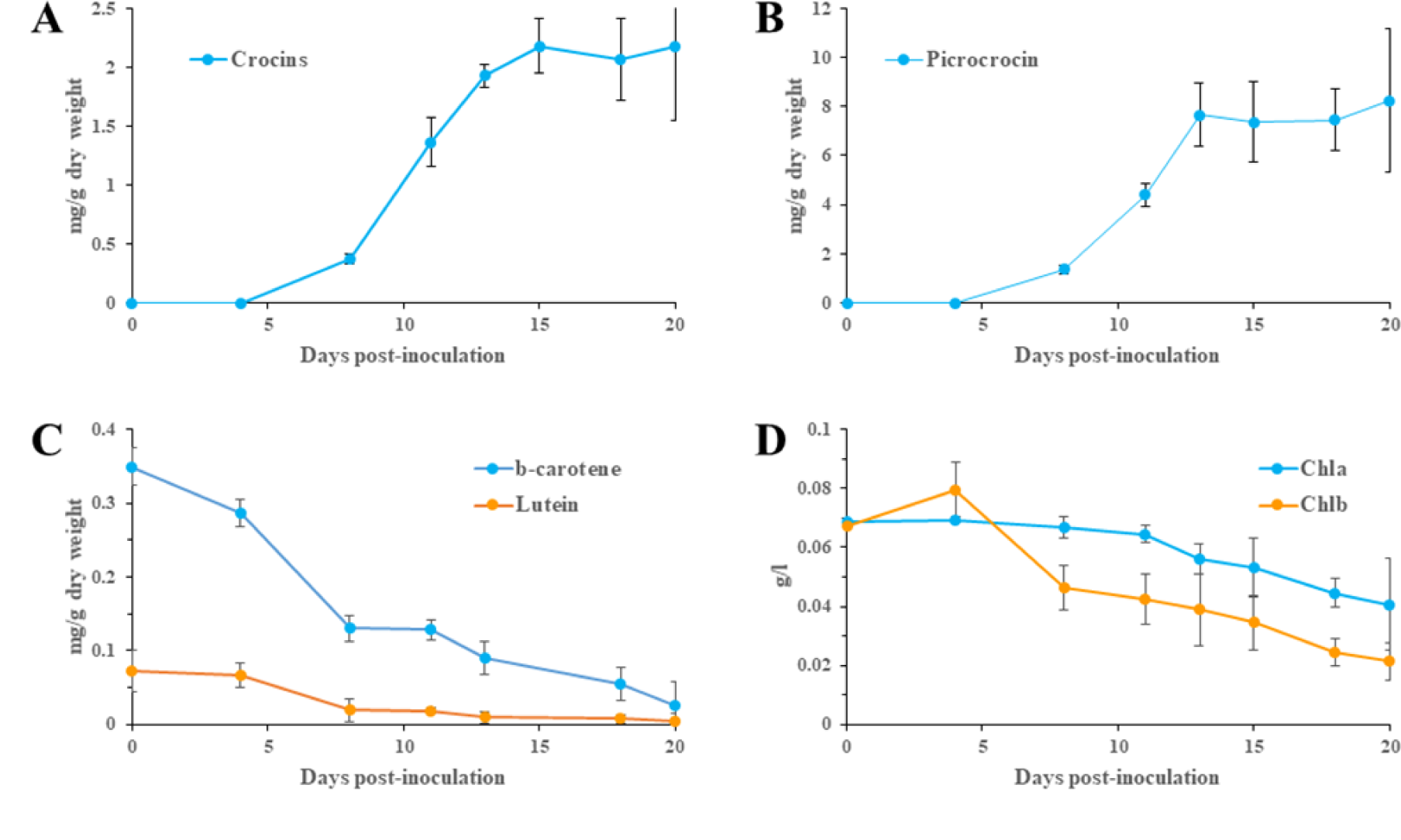
Time-course accumulation of (A) crocins, (B) picrocrocin, (C) β-carotene and lutein, and (D) chlorophyll a and b, as indicated, in systemic tissues of *N. benthamiana* plants agroinoculated with TEVΔNIb-CsCCD2L. Data are average ± sd of at least three biological replicates.

### 3.5. Virus-based combined expression of CsCCD2L and other carotenogenic and non-carotenogenic enzymes

In order to further improve apocarotenoid accumulation in inoculated *N. benthamiana* plants, we combined expression of CsCCD2L with two carotenoid enzymes (Fig. 8A), acting early and late in the biosynthetic pathway, namely *Pantoea ananatis* phytoene synthase (PaCrtB), and saffron β-carotene hydroxylase2 (CsBCH2); or with a non-carotenoid transporter, the saffron lipid transfer protein 1 (CsLTP1), involved in the transport of secondary metabolites (Wang et al., 2016). Interestingly, virus-based co-expression of CsCCD2L and PaCrtB induced a stronger yellow phenotype (Fig. 8B), and a higher content in crocins (3.493±0.325 mg/g DW; Fig. 8C and Table S4). Improved crocins levels were also observed when CsCCD2L was co-expressed with CsBCH2, although at a lesser extent in this case (2.198±0.338 mg/g DW). On the contrary, co-expression of LPT1 did not result in an increment of total apocarotenoid content (Fig. 8C and Table S4). Interestingly, when compared to tissues in which CsCCD2L alone was expressed, a preferential over-accumulation for higher glycosylated crocins was observed, while lower glycosylated species were down-represented (Fig. 8D).

**Fig. 8.**
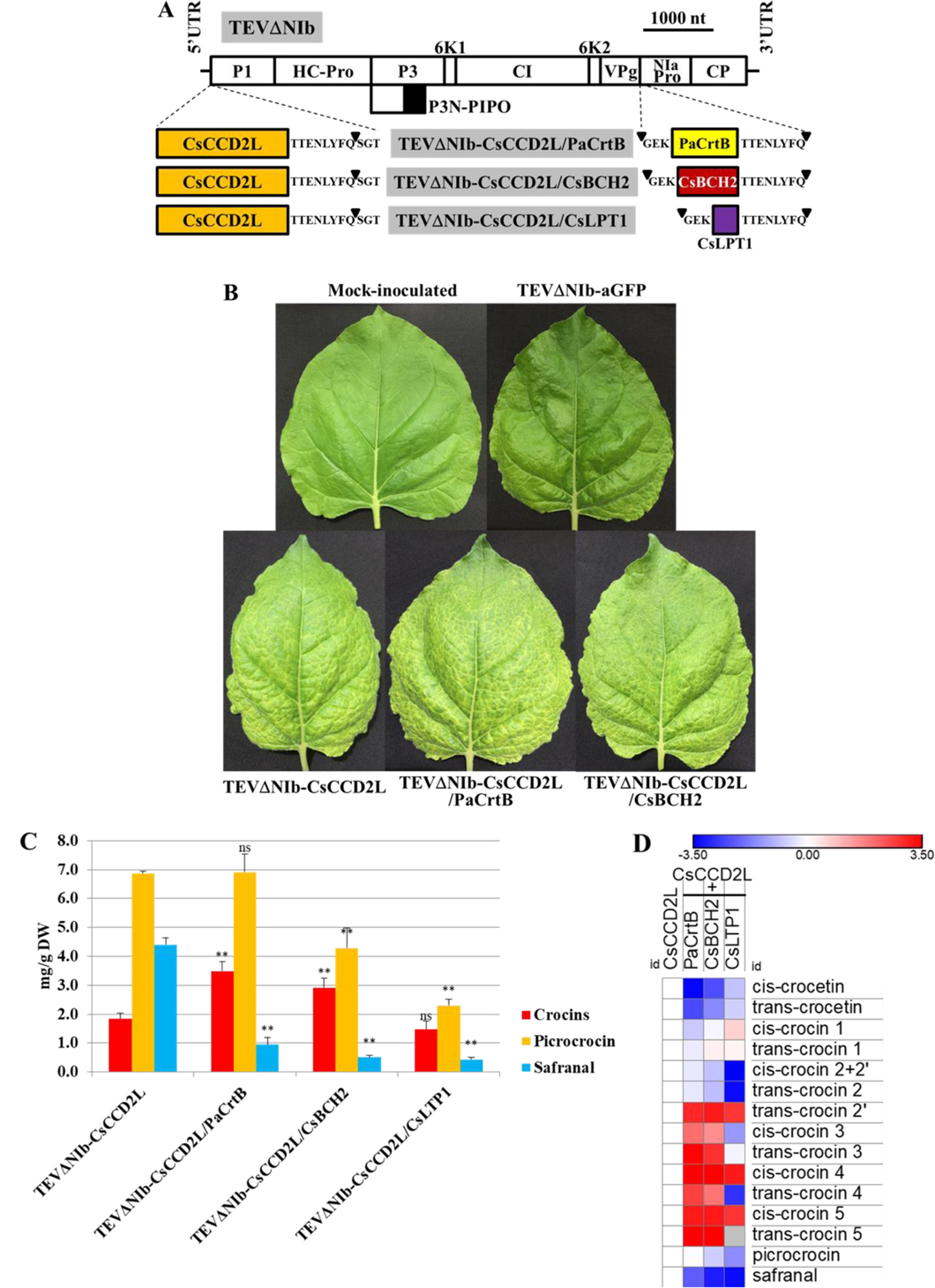
Apocarotenoid accumulation in *N. benthamiana* tissues inoculated with viral vectors carrying CsCCD2L, alone or in combination with PaCrtB, CsBCH2 and CsLTP1. (A) Schematic representation of TEVΔNIb-CsCCD2L/PaCrtB, - CsCCD2L/CsBCH2 and CsCCD2L/CsLPT1. The cDNAs corresponding to CsCCD2L, PaCrtB, CsBCH2 and CsLPT1 are indicated by orange, yellow, red and purple boxes respective. Other details are the same as in Fig. 2A. (B) Representative leaves from plants mock-inoculated and agroinoculated with TEVΔNIb-aGFP, TEVΔNIb-CsCCD2L, TEVΔNIb-CsCCD2L/PaCrtB and TEVΔNIb-CsCCD2L/CsBCH2, at 13 dpi. (C) Total crocins, picrocrocin and safranal contents, expressed as mg/g of DW. Asterisks indicate significant differences according a t-test (pValue: *<0.05; **<0.01; ns, not significant) carried out in the comparisons CsCCD2L vs CsCCD2L+PaCrtB, CsCCD2L+CsBCH2 or CsCCD2L+CsLTP1. (D) Relative accumulation of single crocins, picrocrocin and safranal, normalized on the CsCCD2L-alone sample and visualized as heatmap of log_2_ values. Data are average ± sd of three biological replicates. Analyses were performed at 13 dpi.

## 4. Discussion

The carotenoid biosynthetic pathway in higher plants has been manipulated by genetic engineering, both at the nuclear and transplastomic levels, to increase the general amounts of carotenoids or to produce specific carotenoids of interest, such as ketocarotenoids (Apel and Bock, 2009; Diretto et al., 2019b, 2007; Farré et al., 2014; Giorio et al., 2007; Hasunuma et al., 2008; Nogueira et al., 2019; Zhu et al., 2018). However, these projects also showed that the stable genetic transformation of nuclear or plastid genomes from higher plants is a work-intensive and time-consuming process, which frequently drives to unpredicted results, due to the complex regulation of metabolic pathways. Although no exempted of important limitations, chief among them is the amount of exogenous genetic information plant virus-derived expression systems can carry, they represent an attractive alternative for some plant metabolic engineering goals. Here, we used a virus-driven system that allows the production of noteworthy amounts of appreciated apocarotenoids, such as crocins and picrocrocin, in adult *N. benthamiana* plants in as little as 13 days (Fig. 7). For this, we manipulated the genome of a 10-kb potyvirus, a quick straight-forward process that can be easily scaled-up for the high-throughput analysis of many enzymes and regulatory factors and engineered versions thereof.

In this virus-driven system, the expression of an appropriate CCD in *N. benthamiana* is sufficient to trigger the apocarotenoid pathway in this plant. Recently, it was observed the same situation in a transient expression system (Diretto et al., 2019a). These observations indicate that the zeaxanthin cleavage is the limiting step of apocarotenoid biosynthesis in this species, while aldehyde dehydrogenation and glycosylation steps can be efficiently complemented by endogenous orthologue enzymes. A similar situation was previously observed with other secondary metabolites. Indeed, a large portion of hydroxyl- and carboxyl-containing terpenoid compounds heterologously produced in plants are glycosylated by endogenous UGTs. For example, in transgenic maize expressing a geraniol synthase gene from *Lippia dulcis*, geranoyl-6-*O*-malonyl-β-D-glucopyranoside was the most abundant geraniol-derived compound (Yang et al., 2011). In another study, *N. benthamiana* plants transiently expressing the *Artemesia annua* genes of the artemisin biosynthetic pathway accumulated glycosylated versions of intermediate metabolites (Ting et al., 2013). Likewise, in tomato plants expressing the chrysanthemyl diphosphate synthase gene from *Tanacetum cinerariifolium*, the 62% of trans-chrysanthemic acid was converted into malonyl glycosides (Xu et al., 2018).

Zeaxanthin, the crocins precursor, is not a major carotenoid in plant leaves grown under regular light conditions, although its level increases during high light stress (Demmig-Adams et al., 2012). However, the efficient virus-driven production of all the intermediate and final products of the apocarotenoid biosynthetic pathway in *N. benthamiana* (Fig. 4) suggests that CCD is able to intercept the leaf metabolic flux in order to diverge it towards the production of apocarotenoids. Similarly, the observed reduction in lutein (Fig. 5A) may result from the CCD activity on the β-ring of this molecule, which confirms, at least for the *C. sativus* enzyme, the previous *in vitro* data showing that lutein can act as substrate of this cleavage activity (Frusciante et al., 2014). Although *N. benthamiana* leaves infected with CsCCD2L and BdCCD4.1 recombinant viruses are able to accumulate crocins, it has to be mentioned that, at the qualitative level, their profiles are distinct with respect to the saffron stigma: while in the formers the major crocin species are the trans-crocin 3, together with cis-crocin 4, trans-crocin 2, cis-crocin 3 and cis-crocin 5, saffron stigmas are characterized by the presence, in decreasing order of accumulation, of trans-crocin 4, trans-crocin 3 and trans-crocin 2. These results suggest not only the implication of promiscuous glucosyltransferase enzymes differing form those in saffron, but also the different conditions in which crocins are produce in saffron and *N. benthamiana*, which implies respectively the absence or presence of light during this process, which can effectively influence the cis- or trans-configuration of crocetin (Rubio-Moraga et al., 2010). A higher bioactivity of the trans-forms has been previously demonstrated, at least for crocetin (Zhang et al., 2017), thus making the crocins pool accumulated in the CsCCD2L and BdCCD4.1 leaves a good source of bioactive molecules. The *C. sativus* and *B. davidii* enzymes also generated a different crocins pattern: this finding can be explained by a distinct specificity and kinetics, which might influence the subsequent glucosylation step. Another remarkable observation is the dramatic increase of phytoene and phytofluene levels in *N. benthamiana* tissues as a consequence of the virus-driven expression of CCDs (Fig. 5A). These are upstream intermediates in the carotenoid biosynthetic pathway. Previous reports indicated that the overexpression of downstream enzymes or the increased storage of end-metabolites in the carotenoid pathway can drain the metabolic flux towards the synthesis and accumulation of the end-products (Diretto et al., 2007; Lopez et al., 2008). In our virus-driven system, expression of CsCCD2L and BdCCD4.1 seem to induce the same effect, promoting the activities of the phytoene synthase (PSY) and phytoene desaturase (PDS), which results in a higher accumulation of their metabolic intermediate products phytoene and phytofluene. In this context, the observed amounts of phytoene and phytofluene might be a consequence of PSY and PDS catalyzing the rate-limiting steps of the pathway in *N. benthamiana*, as previously described in other species (Maass et al., 2009; Xu et al., 2006). Levels of additional plastidic metabolites in the groups of tocochromanols and quinones only showed minor fluctuations, which indicates that perturbations in carotenoid and apocarotenoid biosynthesis do not necessarily affect other related pathways that share common precursors, such as geranylgeranyl diphosphate (GGPP) (Fig. 5C). Our results also indicate that the decrease in the chlorophyll levels do not depend on CCD expression, but on viral infection, since it was also observed in the control tissues infected with TEVΔNIb-aGFP (Fig. 5B). This finding agrees with the effect of plant virus infection on photosynthetic machinery (Li et al., 2016).

In our virus-driven experiments, CsCCD2L was more efficient in the production of crocetin dialdehyde and crocins than BdCCD4.1. This suggests that the *C. sativus* CCD used here (CsCCD2L) is definitively a highly active enzyme, completely compatible with the virus vector and the host plant. Indeed, *C. sativus* stigma is the main commercial source for these apocarotenoids (Castillo et al., 2005; Rubio Moraga et al., 2013). Apocarotenoids are also present, although at lower levels, in the flowers of *Buddleja* spp., where they are restricted to the calix.

Virus-driven expression of CsCCD2L in adult *N. benthamiana* plants allowed the accumulation of the notable amounts of 2.18±0.23 mg of crocins and 8.24±2.93 mg of picrocrocin per gram (dry weight) of infected tissues in only 13 dpi (Fig. 7). These amounts are much higher than those obtained with a classic *A. tumefaciens*-mediated transient expression of the CCD2L enzyme in *N. benthamiana* leaves, alone or in combination with UGT709G1 (30.5 μg/g DW of crocins (unpublished data) and 4.13 μg/g DW of picrocrocin (Diretto et al., 2019a), and highlight the opportunity to exploit the viral system for the production of valuable secondary metabolites. A huge range of values has been reported for crocins and picrocrocin in saffron in different studies. Crocins reported values in saffron samples from Spain ranged from 0.85 to 32.40 mg/g dry weight (ALONSO et al., 2001). Other reports showed 24.87 mg/g from China (Tong *et al*., 2015), 26.60 mg/g from Greece (Koulakiotis et al., 2015), 29.00 mg/g from Morocco (Gonda et al., 2012), 32.60 mg/g from Iran, 49.80 mg/g from Italy (Masi et al., 2016) or 89.00 mg/g from Nepal (Li et al., 2018). In *Gardenia* fruits, crocins levels range from 2.60 to 8.37 mg/g (Ouyang et al., 2011; Wu et al., 2014). The picrocrocin content ranges from 7.90 to 129.40 mg/g in Spanish saffron. In Greek saffron, 6.70 mg/g were reported (Koulakiotis et al., 2015), and 10.70-2.16 mg/g in Indian, 21.80-6.15 mg/g in Iranian and 42.20-280.00 mg/g in Moroccan saffron (Lage and Cantrell, 2009). The different values obtained in those samples are also a consequence of the different procedures followed to obtain the saffron spice. Most methods involve a dehydration step by heating, which causes the conversion of picrocrocin to safranal. Although lower than those from natural sources, the crocins and picrocrocin contents of *N. benthamiana* tissues achieved with our virus-driven system may still be attractive for commercial purposes because, in contrast to *C. sativus* stigma or *Gardenia* spp. fruits, *N. benthamiana* infected tissues can be quickly and easily obtained in practically unlimited amounts. Moreover, the virus-driven production of crocins and picrocrocin in *N. benthamiana* may still be optimized by increasing, for instance, the zeaxanthin precursor. PSY over-expression in plants was shown to increase the flux of the whole biosynthetic carotenoid pathway (Nisar et al., 2015), as well as the zeaxanthin pool (Lagarde et al., 2000; Römer et al., 2002). In addition, the virus-based cytosolic expression of a bacterial PSY (*Pantoea ananatis* crtB) also induced general carotenoid accumulation, and particularly increased phytoene levels in tobacco infected tissues (Majer et al., 2017). On the other side, BCH overexpression resulted in increased zeaxanthin and β−β−xanthophyll contents, both in microbes and plants (Arango et al., 2014; Du et al., 2010; Lagarde et al., 2000). In this context, we selected PaCrtB and CsBCH2, together with CsLTP1, involved in carotenoid transport, as additional targets to improve apocarotenoid contents in *N. benthamiana* leaves, which allowed a further increase of 1.9 and 1.6 fold in CsCCD2L/PaCrtB and CsCCD2L/ CsBCH2, respectively, compared to the sole expression of CsCCD2L (Fig. 8). Other targets to improve apocarotenoid accumulation could include the knocking out of enzymatic steps opposing zeaxanthin accumulation, as zeaxanthin epoxydase (ZEP), which uses zeaxanthin as substrate to yield violaxanthin, or lycopene ε-cyclase (LCY-ε) that converts lycopene in δ-carotene, thus diverging the metabolic flux towards the synthesis of ε−β− (lutein), rather than β−β−xanthophylls (as zeaxanthin), which will be object of future attempts.

In summary, here we describe a new technology that allows production of remarkable amounts of highly appreciated crocins and picrocrocin in adult *N. benthamiana* plants in as little as 13 dpi. This achievement is the consequence of *N. benthamiana*, a species that accumulate negligible amounts of apocarotenoids, only requiring the expression of an appropriate CCD to trigger the apocarotenoid biosynthetic pathway, and can be further improved through the combination with the overexpression of early or late carotenoid structural genes.

## Acknowledgements

We thank K. Schreiber and C. Mares (IBMCP, CSIC-UPV, Valencia, Spain) for technical assistance during plant transformation. We thank M. Gascón and M.D. Gómez-Jiménez (IBMCP, CSIC-UPV, Valencia, Spain) for helpful assistance with LSCM analyses. We thank D. Dubbala (IBMCP, CSIC-UPV, Valencia, Spain) for English revision. This work was supported by grants BIO2016-77000-R and BIO2017-83184-R from the Spanish Ministerio de Ciencia, Innovación y Universidades (co-financed European Union FEDER funds). M.M. was the recipient of a predoctoral fellowship from the Spanish Ministerio de Educación, Cultura y Deporte (FPU16/05294). G.D. and L.G.G. are participants of the European COST action CA15136 (EUROCAROTEN). L.G.G. is a participant of the CARNET network (BIO2015-71703-REDT and BIO2017-90877-RED).

---

**Fig. S1.**
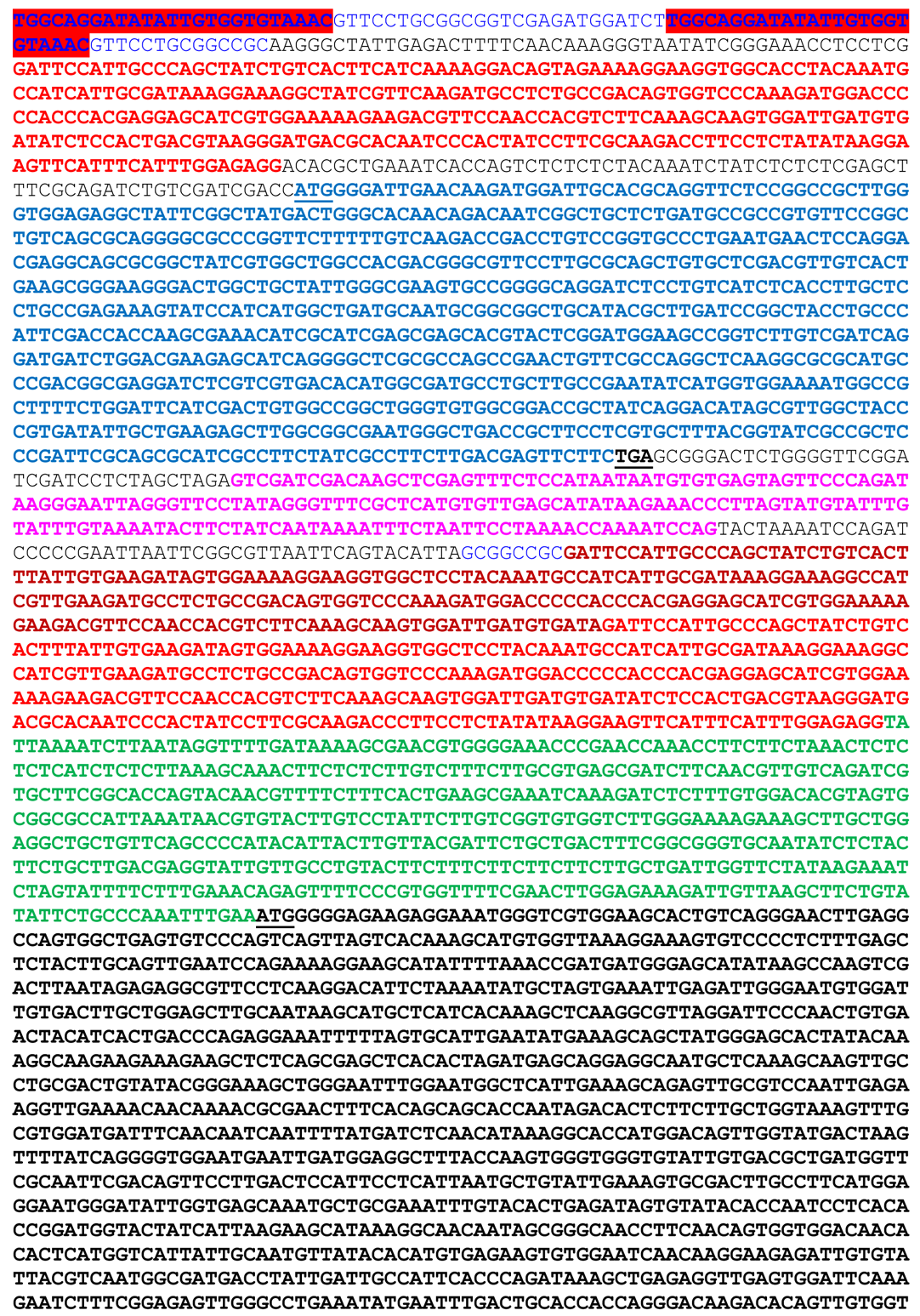
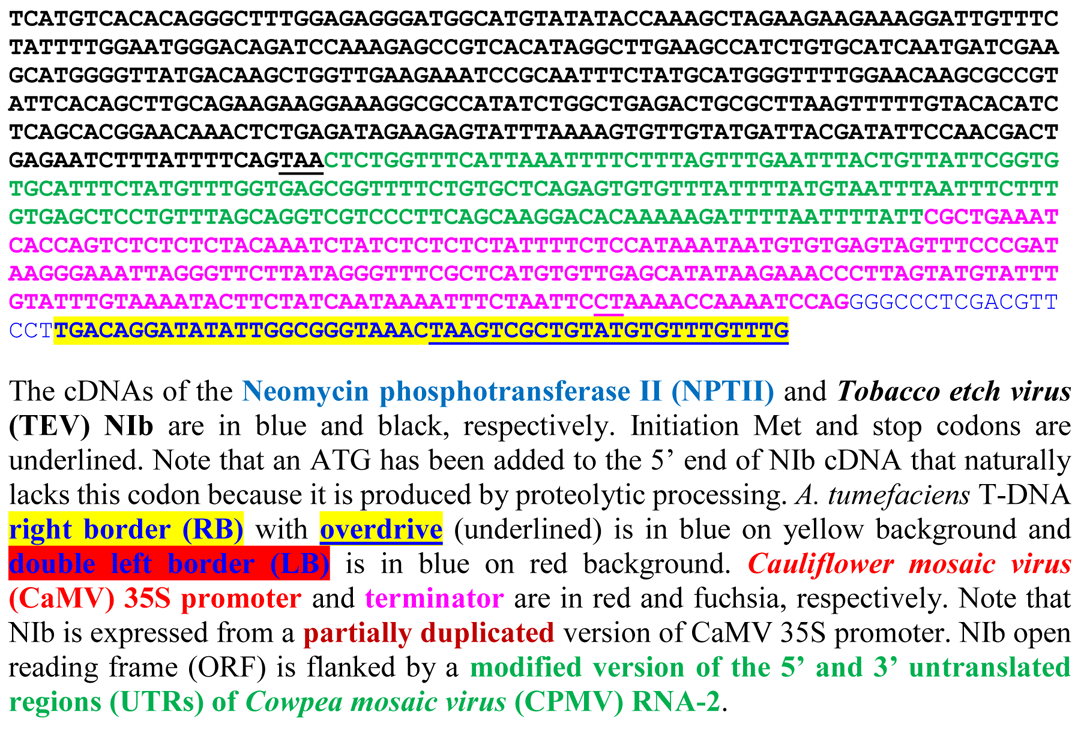
Nucleotide sequence of the cDNA transferred to the *Nicotiana benthamiana* transformed line using *Agrobacterium tume*faciens to prepare the virus-drive expression system.

**Fig. S2.**
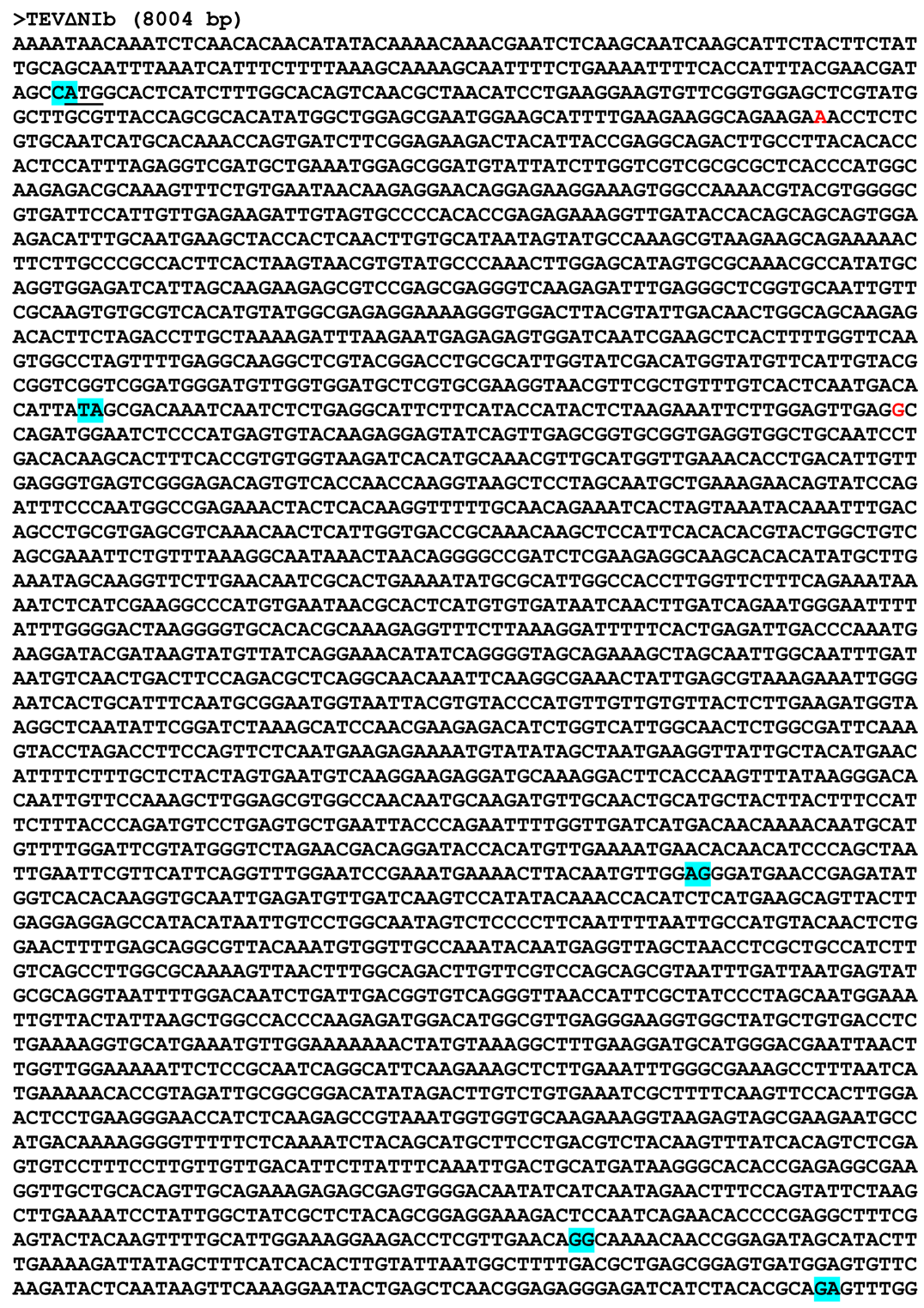
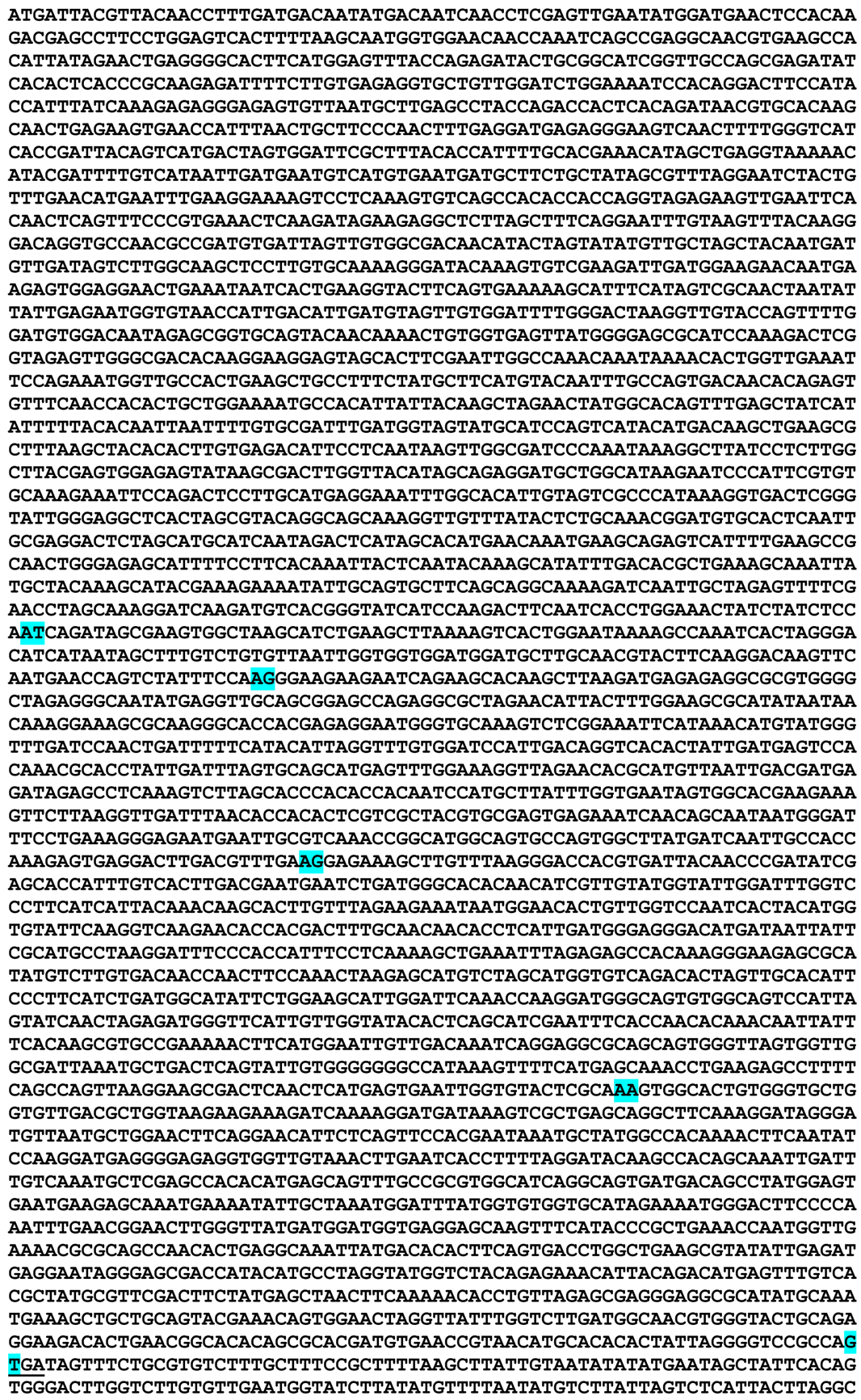
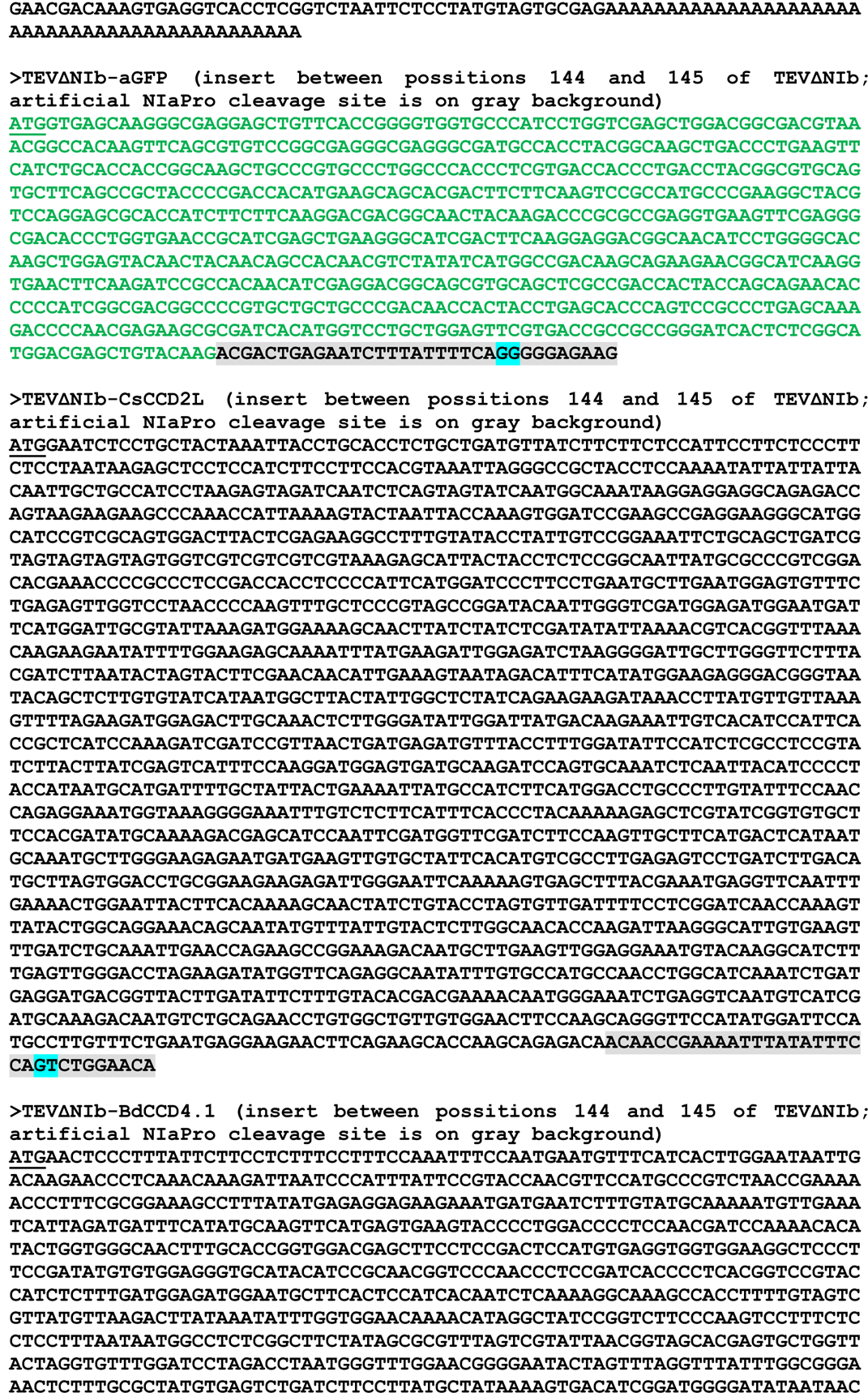
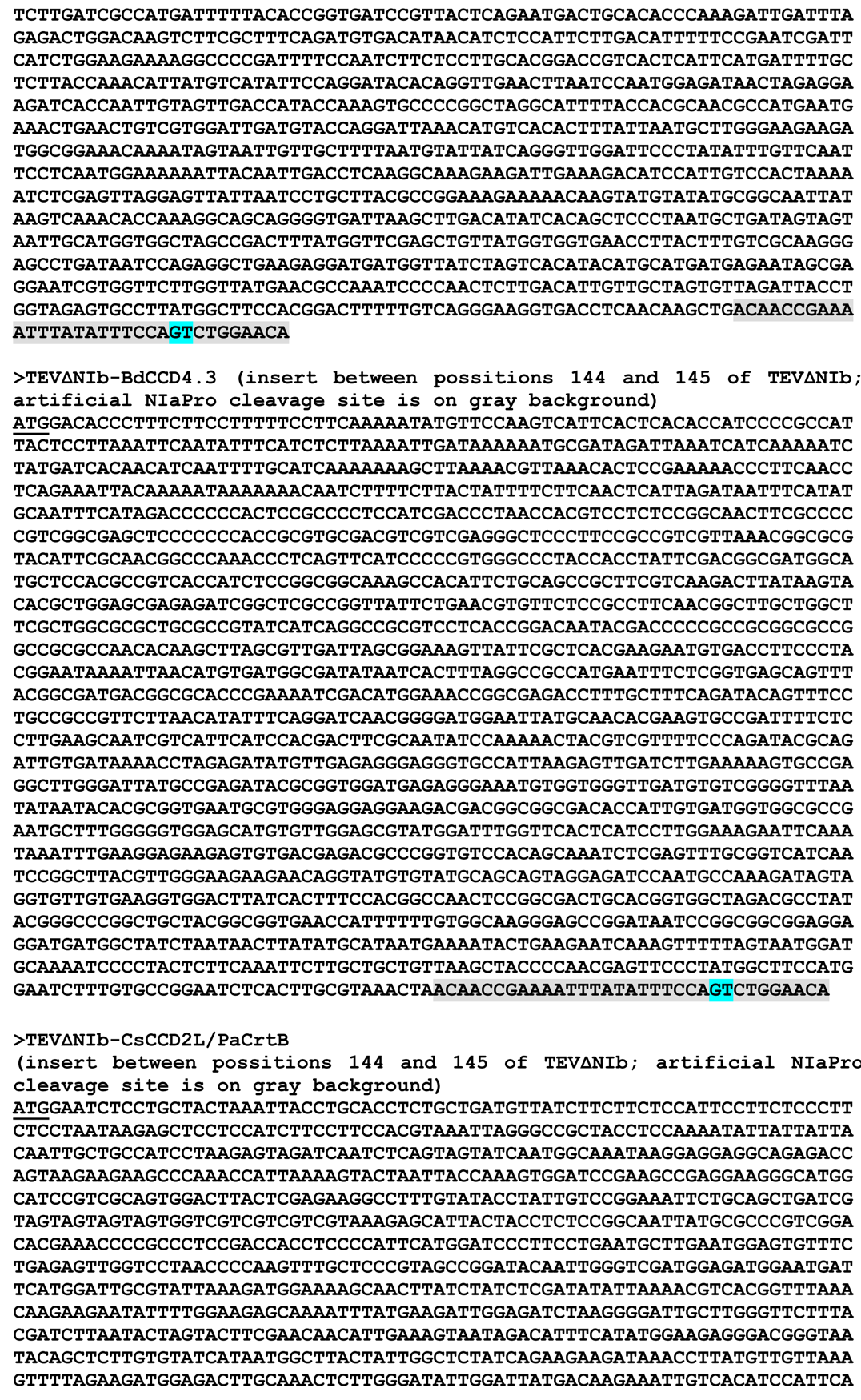
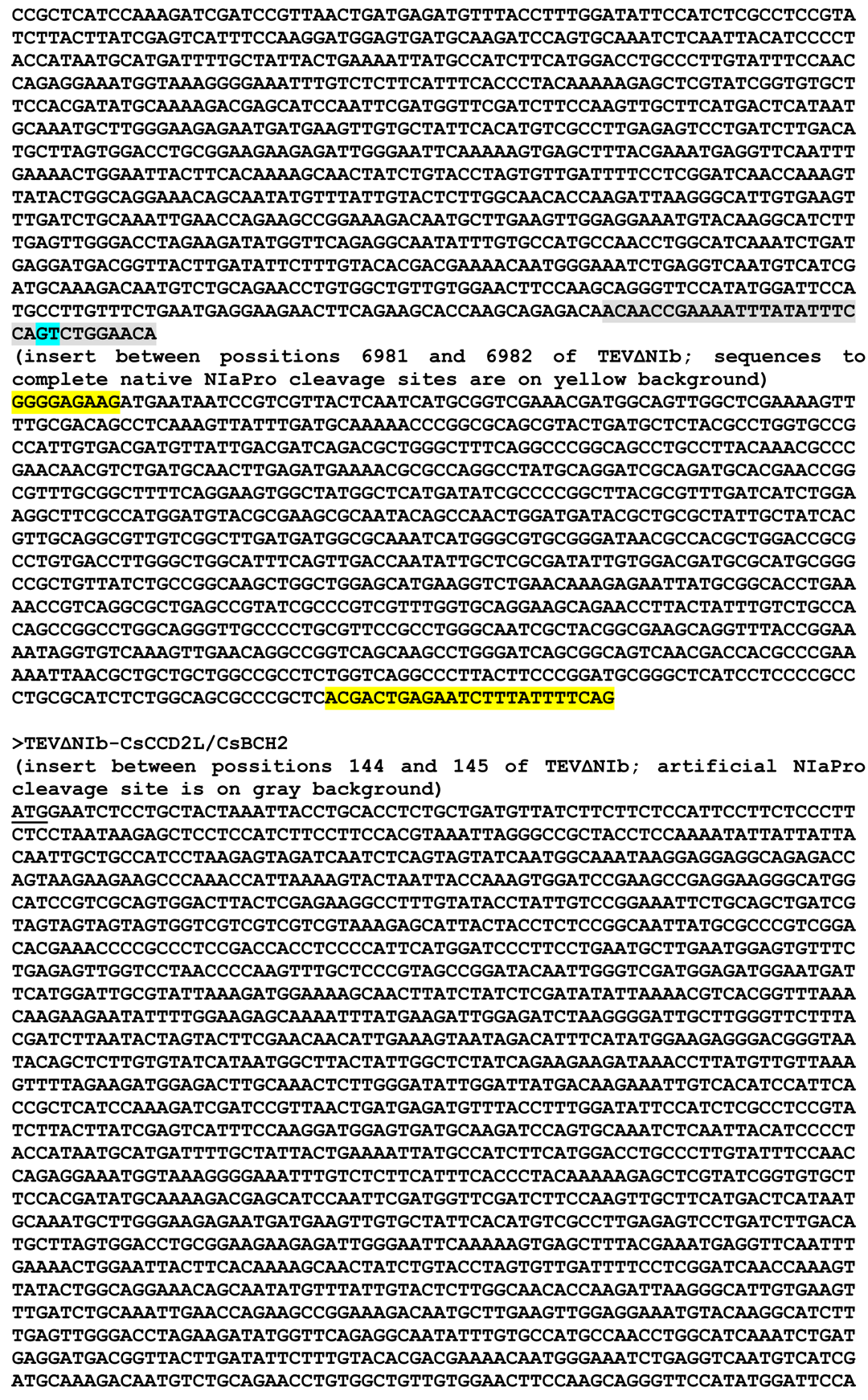
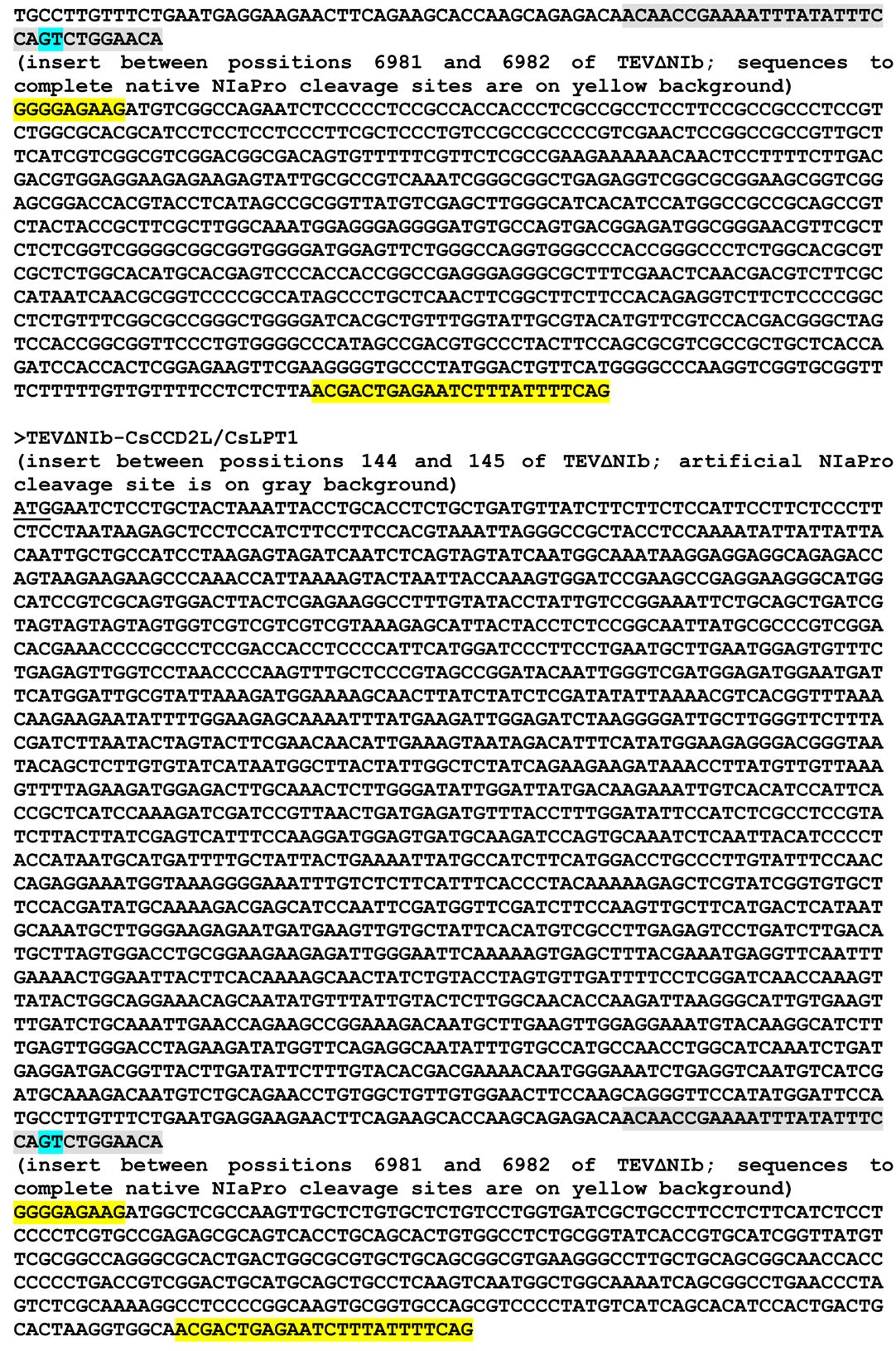
Nucleotide sequences of TEVΔNIb (sequence variant DQ986288 that include the two silent mutations G273A and A1119G, in red, and lacks the whole NIb cistron) and the derived recombinant viruses TEVΔNIb-aGFP, -CsCCD2L, -BdCCD4.1, -BdCCD4.3, -CsCCD2L/PaCrtB, -CsCCD2L/CsBCH2 and -CsCCD2L/CsLPT1. The boundaries of viral cistrons are indicated on blue background. Initiation Met and stop codons are underlined.

**Fig. S3.**
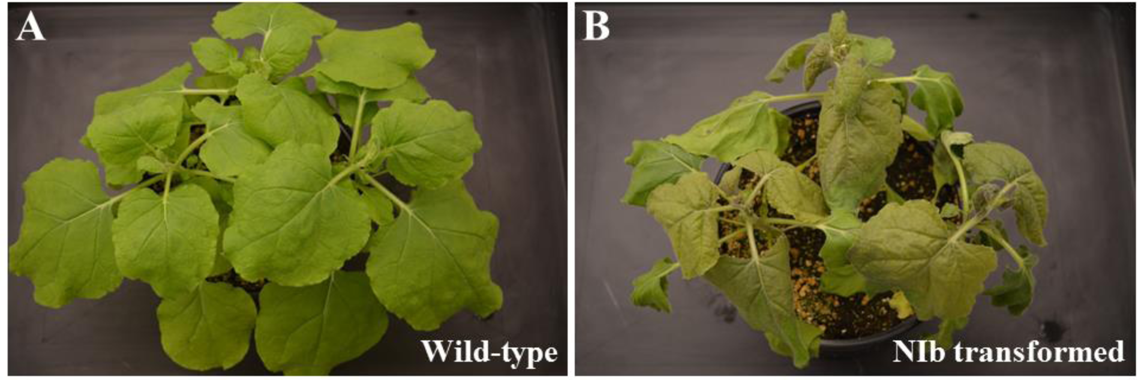
Complementation of a TEV mutant that lacks NIb in the transformed *N. benthamiana* line that stably expresses TEV NIb. (A) Wild-type and (B) NIb-expressing *N. benthamiana* plants were mechanically inoculated with TEVΔNIb-Ros1, a recombinant virus in which the NIb cistron is replaced by the Rosea1 visual marker that induces anthocyanin accumulation. Pictures were taken 11 days post-inoculation (dpi).

**Fig. S4.**
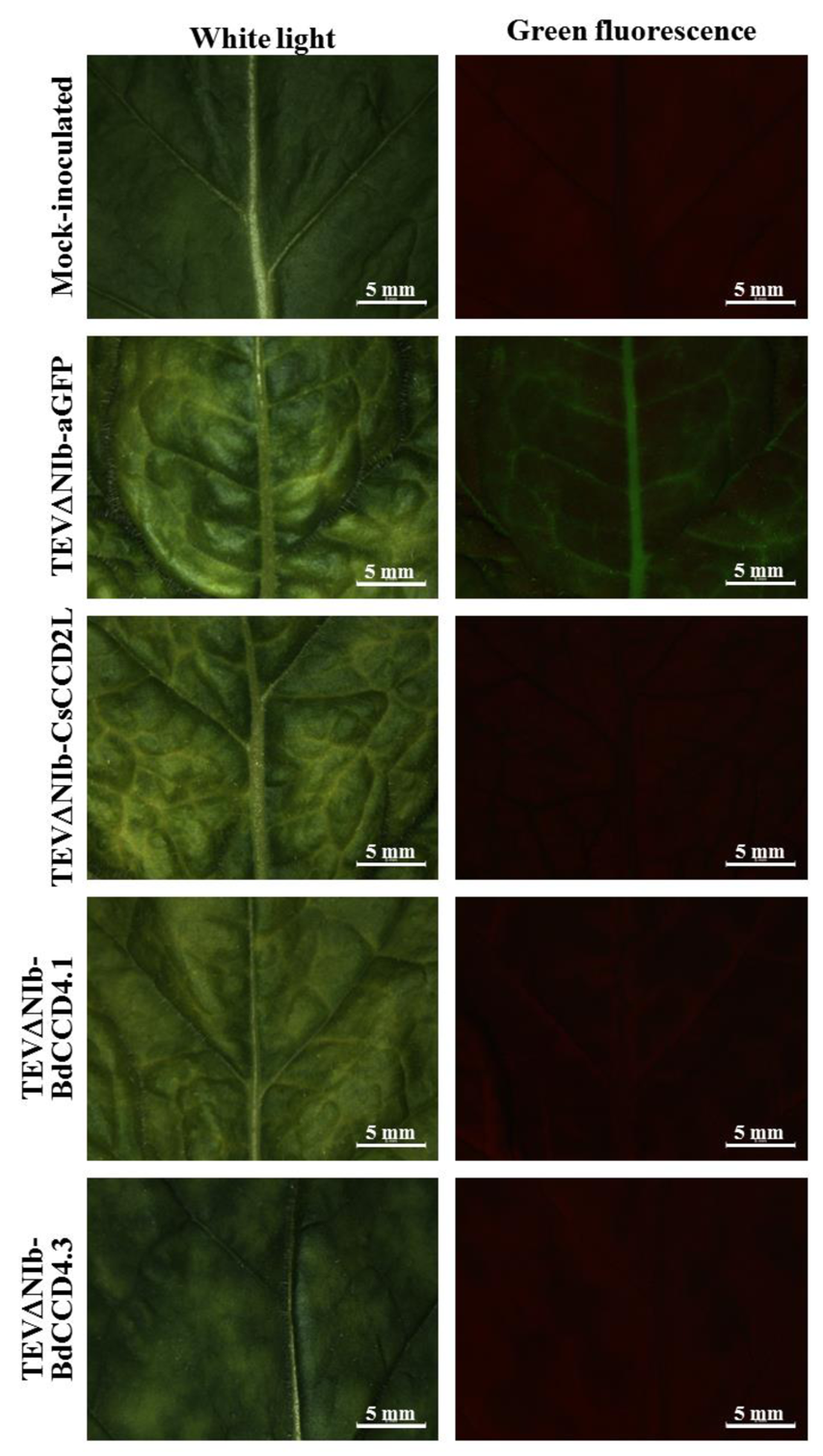
Fluorescence stereomicroscope analysis of tissues from plants mock-inoculated and agroinoculated with TEVΔNIb-aGFP, TEVΔNIb-CsCCD2L, -BdCCD4.1 and - BdCCD4.3. Pictures at 15 dpi were taken with a Leica MZ 16 F stereomicroscope under white light and under UV illumination with the GFP2 (Leica) fluorescence filter. Scale bars correspond to 5 mm.

**Fig. S5.**
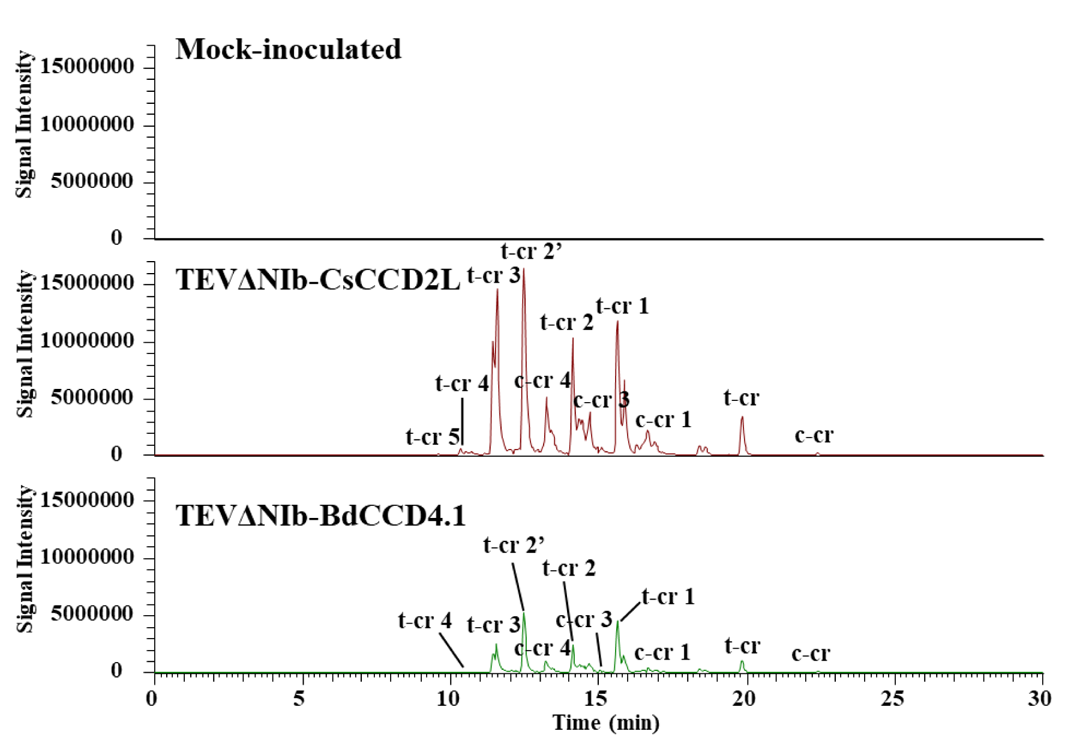
Crocetin ion extraction (m/z 329.1747 (M+H+)) in control and CsCCD2L and BdCCD4.1 polar fraction chromatograms. Metabolites were extracted from infected leaves and run on an HPLC-PDA-HRMS system. The apocarotenoids have an accurate mass and a chromatographic mobility identical to those of the authentic standards, when available.

**Table S1.** LC-HRMS of crocins-(A) and picrocrocin-related (B) apocarotenoid levels in infected leaves. Data are avg ± sd of, at least, three biological replicates; nd (not detected). Values are reported as fold level between the signal intensities of the apocarotenoid metabolite on the signal intensity of the internal standard (formononetin).

**Table S2.** Relative abundance, as % on the total, of apocarotenoid levels in CsCCD2L and BdCCD4.1 leaves. (A) crocins-related metabolites and (B) picrocrocin-related metabolites.

**Table S3.** LC-HRMS of carotenoid-(A), chlorophyll-related (B), and tocochromanols and quinones (C) levels in infected leaves. Data are avg ± sd of at least three biological replicates; nd (not detected). Values are reported as fold level between the signal intensities of the apocarotenoid metabolite on the signal intensity of the internal standard (a-tocopherol acetate). Asterisks indicate significant differences according a t-test (pValue: *<0.05; **<0.01) carried out in the comparisons Not infected vs GFP, CCD2L vs GFP and CCD4.1 vs GFP.

**Table S4.** LC-HRMS of crocins-(A) and picrocrocin and safranal (B) apocarotenoid levels in infected leaves. Data are avg ± sd of at least three biological replicates; nd (not detected). Values are reported as absolute amounts (in mg/g DW) by using authentical standards and external calibration curves. Asterisks indicate significant differences according a t-test (pValue: *<0.05; **<0.01) carried out in the comparisons CsCCD2L vs CsCCD2L+PaCrtB, CsCCD2L+CsBCH2 or CsCCD2L+CsLTP1.

